# Mechanistic Genome Folding at Scale through the Differentiable Loop Extrusion Model

**DOI:** 10.1101/2025.10.17.682904

**Authors:** Tina Subic, Ali Tuğrul Balcı, Kristina Perevoshchikova, Diego Borges-Rivera, Jieni Hu, Geoffrey Fudenberg, Jacqueline Dresch, Maria Chikina

## Abstract

The spatial folding of the genome shapes gene regulation by controlling which loci interact, yet inferring the mechanisms behind these 3D structures from contact maps remains difficult. Cohesin-mediated loop extrusion is a key organizer of domains and loops, but existing methods either predict contacts without mechanistic insight or simulate extrusion with limited scalability.

We present the differentiable loop extrusion model (dLEM), a scalable framework that reformulates extrusion as a smooth, trainable process. dLEM represents extrusion through position-specific velocity profiles for leftward and rightward cohesin movement. Fitting dLEM to chromosome conformation capture data yields a one-dimensional, interpretable description of extrusion dynamics that aligns with genomic and epigenomic features. dLEM parameters also capture architectural changes under CTCF and WAPL perturbations, enabling genome-wide prediction of extrusion disruptions.

Extending our observations, we demonstrate that dLEM can be seamlessly incorporated into deep learning models to infer extrusion parameters directly from sequence and chromatin features, reducing model complexity by nearly three orders of magnitude while preserving predictive accuracy. Indeed, when incorporated into deep dLEM, dLEM acts as a biophysically-motivated layer for long range genomic communication, and together they provide a predictive, interpretable framework linking 1D genomic features to 3D chromatin folding and its response to sequence and chromatin state, with dLEM’s mechanistic modeling enabling prediction of trans factor perturbation effects.

## Introduction

The genome’s three-dimensional organization adds a regulatory layer beyond sequence, bringing distant loci into proximity to control gene expression and cell identity. Disruptions in folding are linked to developmental disorders and disease, underscoring its central role in biology [1]. Chromosome conformation capture technologies such as Hi-C and Micro-C [2] have revealed a hierarchical structure: at megabase scales, active and inactive compartments; at intermediate scales, topologically associating domains (TADs) and chromatin loops often anchored by CTCF; and at finer scales, enhancer–promoter hubs essential for precise transcriptional control [3–6]. These features vary across cell types and developmental stages, linking genome architecture directly to cellular function [7].

Loop extrusion has emerged as a central mechanism organizing genomes. In this process, cohesin translocates along chromatin, progressively extruding loops until blocked by boundary elements, typically CTCF bound in convergent orientation. Elegant in vitro and in vivo studies explain characteristic contact map patterns—stripes, loops, and domains—through this mechanism [8–11] (Fig. 1A). Yet CTCF alone cannot account for the full complexity of chromatin folding. Transcriptional activity and associated chromatin features also modulate cohesin dynamics [12, 13], suggesting a pervasive yet imperfectly quantified interplay between extrusion and gene regulation [14–16].

**Fig. 1:**
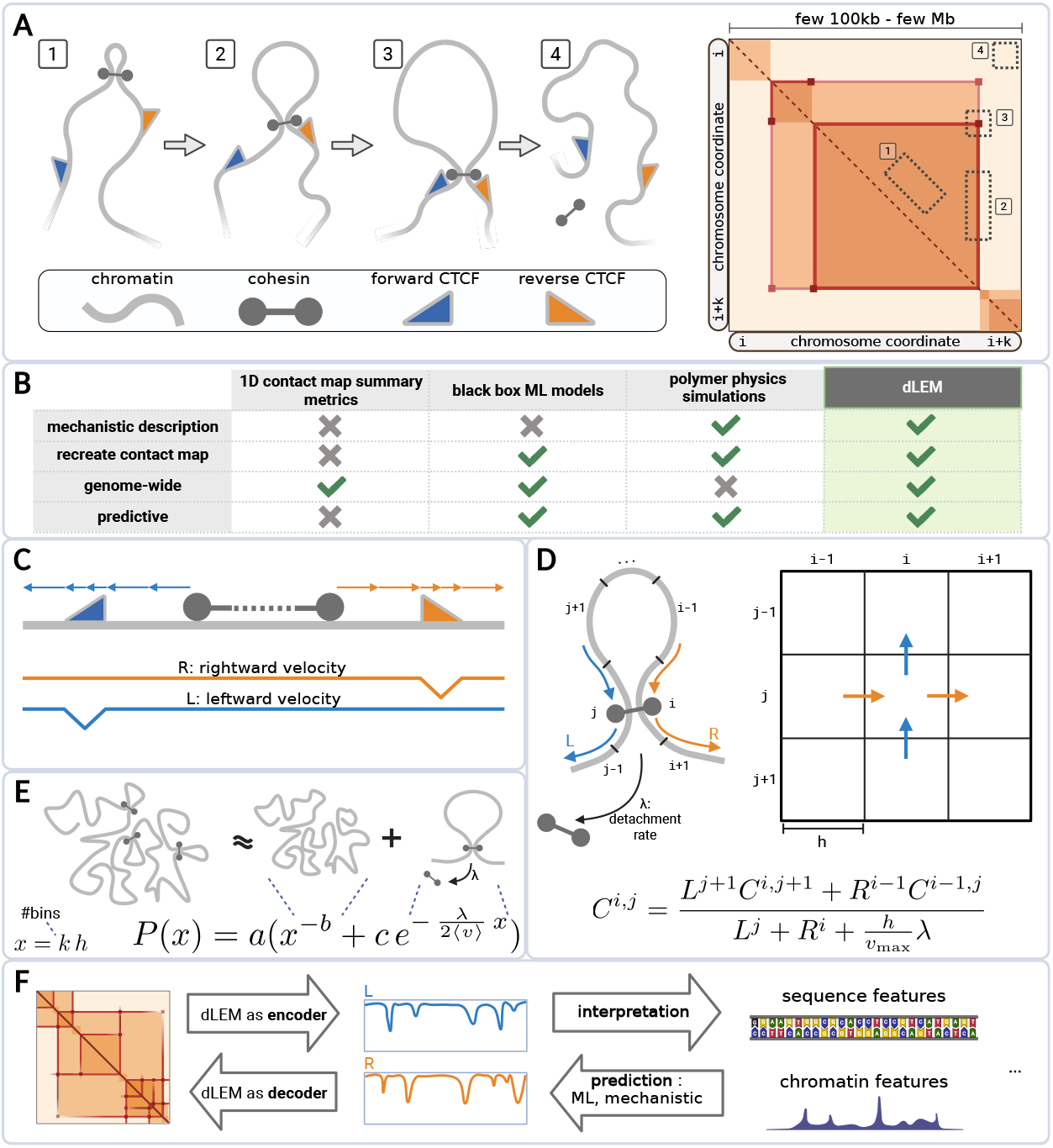
**A**: *Schematic of loop extrusion and resulting contact map features. 1* : Cohesin (grey barbell) binds chromatin and translocates bidirectionally, progressively bringing distant loci into contact (dark regions away from the diagonal). *2* : CTCF proteins bound in forward (blue) or reverse (orange) orientation block cohesin movement directionally. When one cohesin leg encounters CTCF, continued movement of the opposite leg creates asymmetric contact patterns visible as stripes. *3* : When both legs encounter convergent CTCF sites, stable loops form with high anchor contact frequency (dark dots). The intervening region forms a Topologically Associating Domain (TAD) with elevated internal contacts. *4* : Cohesin release by WAPL limits translocation distances. **B**:*Comparison of computational approaches for contact map analysis*. dLEM combines advantages of existing methods while satisfying all criteria for mechanistic loop extrusion analysis. **C**: *dLEM parameterization of loop extrusion dynamics*. Cohesin translocation is decomposed into independent leftward (*L*) and rightward (*R*) components, each with position-specific velocities. Obstacles such as CTCF binding sites reduce translocation speed, creating dips in the corresponding velocity profiles. **D**: *Mathematical formulation of dLEM*. The steady-state equation (shown) describes how contact frequency *C*^*i,j*^ between loci *i* and *j* arises from steady state of cohesin flux. Cohesin flows in from preceding neighboring contacts (numerator terms) and flows out through continued translocation and detachment at rate *λ* (denominator). The bin resolution *h* maps molecular-scale dynamics to chromosome conformation capture experimental resolution, directly connecting chromatin fragment transitions to contact map pixel transitions. **E**: *Distance-decay analysis reveals cohesin detachment rate*. Average contact frequency as a function of genomic distance *x* combines two independent physical processes: polymer folding (power-law decay, exponent *−b*) and loop extrusion (exponential decay, rate 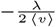). The model treats these as additive contributions, where *c* represents the loop extrusion contribution and *a* provides data scaling. This physics-based approach directly measures the detachment rate parameter *λ*. **F** *dLEM versatility*: dLEM functions as both encoder and decoder: (1) encoding Hi-C contact maps into 1D parameters for comparison with sequence features, chromatin tracks, or trans factor perturbations; (2) predicting contact maps from genomic features via mechanistic or machine learning models that generate dLEM parameters and decode them into contact maps.

A key barrier to understanding this relationship is the dimensional mismatch between data types: chromosome conformation capture experiments yield twodimensional contact matrices, while sequence and chromatin features are inherently one-dimensional. This mismatch has made it difficult to systematically quantify how CTCF, transcriptional machinery, and other chromatin and sequence features modulate loop extrusion dynamics, and to quantitatively predict how genetic or molecular perturbations would reshape chromatin architecture. Current computational strategies face fundamental trade-offs that prevent them from bridging the dimensional gap in a mechanistic and interpretable way: summary statistics quantify patterns like insulation but lack mechanistic grounding [5, 6, 17, 18]; polymer physics simulations are mechanistically rich but not scalable genome-wide [8, 9, 19]; and deep learning models such as DeepC [20], Akita [21], Orca [22], and C.Origami [23] can predict contact maps at scale (for review, see [24]), but act as black boxes, making it difficult to extract mechanistic insight or relate predictions directly to physical processes such as loop extrusion. The fundamental challenge is to bridge this dimensional gap through a framework that is simultaneously mechanistic, interpretable, and scalable.

To overcome these limitations (Fig. 1B), we developed the differentiable loop extrusion model (dLEM), which reformulates loop extrusion as a genome-wide differentiable process. dLEM solves the dimensional mismatch by reducing complex 2D contact maps into simple 1D extrusion parameters. Specifically, dLEM parameterizes extrusion through two position-specific velocity profiles—leftward (L) and rightward (R)—that capture how local genomic context modulates cohesin translocation speed at each position. Obstacles such as CTCF binding sites appear as characteristic dips in these velocity profiles (Fig.1C). These simple 1D patterns generate the complex 2D structures observed in contact maps: TADs form between boundaries with convergent orientation (low *L* downstream, low *R* upstream); stripes emerge from asymmetric blocking where only *L* or only *R* is low; and focal loops appear where cohesin stalls at paired positions with low *L* and *R*. Optimizing these velocity profiles against contact map data yields a one-dimensional representation of extrusion dynamics that aligns naturally with genomic tracks, bridging descriptive correlation and predictive mechanistic modeling. Critically, because these parameters directly reflect physically interpretable extrusion dynamics, they can be perturbed in silico, enabling quantitative predictions of structural outcomes under experimental manipulations of loop extrusion components, such as CTCF or WAPL depletion.

Building on this framework, we develop deep dLEM, which enables prediction of contact maps across cellular contexts from sequence and widely-available chromatin accessibility profiles. deep dLEM learns to predict the slowdown profiles directly from genomic sequence and chromatin features, and then generates contact maps through the mechanistic dLEM process. In this sense, dLEM can be viewed as the first biophysically-motivated differentiable layer for long-range information propagation in the genome, seamlessly compatible with the broader deep learning “stack” for genomics. Deep dLEM simply combines this layer with upstream sequence and chromatin encoders, constraining learning to the same physically grounded latent space. This design preserves interpretability while demonstrating that fully predictive modeling of genome architecture from sequence and one-dimensional genomic data can be achieved within a mechanistic, physically meaningful framework.

Together, dLEM and its deep learning application (deep dLEM) shift the focus from predicting contact map patterns to uncovering the biophysical dynamics that generate them (Fig. 1F). By connecting one-dimensional genomic landscapes to threedimensional chromatin architecture through interpretable parameters, our framework not only explains observed variation but also predicts structural responses to perturbations, opening avenues for decoding developmental regulation, interpreting disease-associated variants, and ultimately engineering genome organization.

## Results

### Complex contact maps emerge from simple one-dimensional dynamics

**differentiable loop extrusion model (dLEM)** bridges the dimensional gap between 2D contact maps and 1D genomic features by decomposing contact matrices into interpretable extrusion parameters. The key insight is that loop extrusion, though it generates complex 2D contact patterns, operates through simple 1D dynamics: cohesin binds and translocates along the linear chromatin sequence (Fig. 1A). Where cohesin slows (at bound CTCF sites, active transcription, and other chromatin features) creates position-specific velocity patterns along the genome. It is these 1D velocity patterns that generate the 2D structures observed in contact maps: TADs form between convergent boundaries, stripes emerge from asymmetric blocking, and loops appear at stalled positions. dLEM recovers these underlying 1D parameters from observed 2D contact data, enabling direct mechanistic interpretation and quantitative prediction.

Inspired by the stochastic 1D simulation of loop-extruding 1D simulation of loop-extruding factors (LEF) dynamics [8], we parameterized loop extrusion through two one-dimensional velocity profiles, leftward (*L*) and rightward (*R*), that capture how fast cohesin moves at each genomic position, with impediments appearing as characteristic dips in these velocity landscapes (Fig. 1C) and causing cohesins to accumulate and form loops. At each genomic position pair (*i, j*), cohesin flows in from the preceding neighboring pixels, and flows out due to continued movement to the succeeding neighboring pixels, or detachment (Fig. 1D). Using the two velocity parameters and the detachment rate, we reconstruct contact maps by modeling the steady-state flow of cohesin density at each pixel, where contact frequency reflects the equilibrium balance of cohesin influx and outflux.

While this formulation makes simplifying assumptions (treating *λ* as constant rather than position-specific and neglecting cohesin-cohesin interactions), these approximations enable tractable genome-wide inference while capturing the essential physics (detailed assumptions in Appendix 5). To fully specify the model, we incorporated a distance-decay analysis that captures how average contact frequency decreases with genomic separation through two distinct physical processes: polymer folding (power-law decay [25]) and loop extrusion (exponential decay controlled by *λ*) (Fig. 1E). This complete framework—local velocity parameters plus global detachment rate—provides a mechanistic description of how 1D dynamics generate 2D contact patterns, with the key innovation being its differentiable formulation that enables efficient genome-wide optimization using modern automatic differentiation frameworks.

When dLEM is fit directly to experimental Micro-C data, it acts as an encoder–decoder: the model compresses 2D contact maps into two 1D velocity profiles and a single detachment rate parameter, and then reconstructs the map from these parameters. As shown on a representative genomic region, the velocity impediments of the fitted parameters directly generate the observed nested TADs, loops, and stripes in the reconstructed contact map and the reconstruction is highly accurate with median correlation *r* = 0.68 in observed over expected quantification (Fig. 2A).

**Fig. 2:**
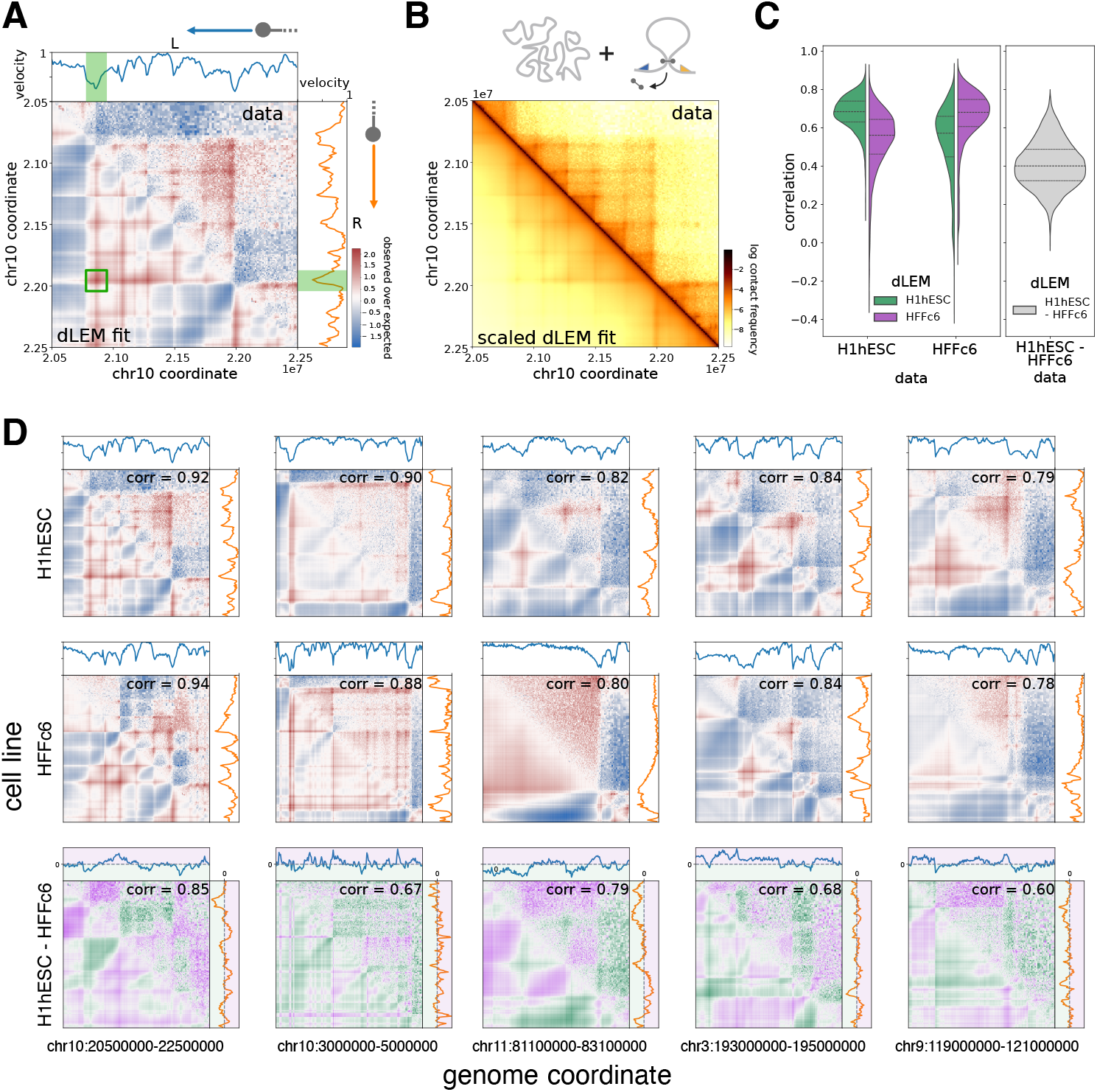
dLEM extracts cell-type specific differences form Micro-C contact maps. **A:** *Example dLEM fit to Micro-C contact map*. Representative 2Mb genomic region (chr10:20.5-22.5Mb) of H1hESC at 10kb resolution showing observed-over-expected experimental data (upper triangle) and dLEM model fit (lower triangle). Red indicates enriched contacts, blue indicates depleted contacts relative to expected values. Fitted leftward (L, blue, top) and rightward (R, orange, right) velocity parameters are shown alongside corresponding map dimensions. The velocity dips highlighted in green result in a loop highlighted with a green box. **B**: *dLEM performance on native contact frequency scale*. Same region as Panel A at 10kb resolution displaying log contact frequencies (darker colors indicate higher contact frequency) with experimental data (upper triangle) and dLEM prediction scaled by incorporating diffusion-extrusion model terms (lower triangle). **C**: *Genome-wide validation of dLEM performance across cell types*. Violin plots showing correlations between dLEM fits and experimental data across sliding window patches from chromosomes 3-14. Left panel compares H1hESC (green) and HFFc6 (purple) model performance, demonstrating higher correlations when models are matched to their corresponding cell types. Right panel shows correlation between experimental cell line differences and dLEM-predicted differences, confirming that dLEM accurately captures cell type-specific chromatin architecture genome-wide. **D**: *Representative examples of cell type-specific chromatin architecture*. Five representative 2Mb genomic regions comparing H1hESC (top row) and HFFc6 (middle row) cell lines, with experimental data (upper triangles) and dLEM fits (lower triangles). Fitted velocity parameters L and R are shown alongside each contact map. Cell line differences (bottom row) show regions where H1hESC has higher contact frequency (green) or HFFc6 has higher contact frequency (purple) in both contact maps and velocity parameters (green/purple shaded regions in L and R plots are denoting greater slowdown in H1hESC/HFFc6), demonstrating dLEM’s ability to recapitulate cell type-specific chromatin architecture.

Furthermore, by incorporating the distance-decay model (Fig. S1), dLEM reproduces not just relative contact patterns but actual contact frequencies on the native scale (Fig. 2B). The performance remains robust across resolutions: even at 2kb resolution where experimental noise increases substantially, the inferred velocity parameters remain consistent with those obtained at 10kb, demonstrating that we are capturing genuine biological signal (Fig. S2).

This biological signal extends to cell type-specific differences: genome-wide analysis shows that models fitted to one cell type show reduced performance when applied to another, indicating that the velocity parameters encode meaningful biological variation (Fig. 2C). The model successfully reconstructs diverse chromatin architectures across the genome and captures cell type-specific differences: when fitted separately to human embryonic stem cell and fibroblast Micro-C data, dLEM recovers distinct velocity landscapes that accurately predict where each cell type shows enhanced or depleted contacts (Fig. 2D). These consistent, high-performance fits across scales and cell types demonstrate that complex chromatin architecture indeed emerges from simple one-dimensional dynamics.

### Transcriptional machinery shapes cohesin dynamics beyond CTCF

Loop extrusion is classically understood as being governed by CTCF boundaries: convergent motifs stall cohesin translocation and generate the asymmetric stripes and focal loops seen in contact maps. This paradigm explains many architectural features, but it leaves open how much other chromatin-associated processes contribute at the genome scale. A key advantage of dLEM is that its one-dimensional extrusion parameters can be directly compared with genomic annotations, enabling systematic dissection of the determinants of cohesin slowdown.

To validate dLEM’s biological relevance and enable systematic comparison with genomic annotations, we first benchmarked fitted velocity parameters against diamond insulation scores, a widely used metric for quantifying TAD boundaries from contact maps (Fig. S3). min(*L, R*) correlates strongly with insulation scores, yet it identifies CTCF-bound boundaries more effectively (Fig. S3 C, D). Indeed, the directional *L* and *R* parameters reveal biology that insulation scores cannot capture: *L* is significantly lower at forward-oriented CTCF sites while *R* is lower at reverse-oriented sites, whereas insulation scores show no directional preference (Fig. S3 E, F). This suggests that dLEM deconvolves chromatin architecture into its directional components.

Indeed, across three cell types (H1hESC, IMR90, and K562), fitted velocity parameters consistently correlated most strongly with the extrusion machinery itself (RAD21, CTCF, and directional CTCF motifs) as well as with open chromatin (DNase-seq) (Fig. 3A). Interestingly, CTCF motifs themselves had a strong preference for either the leftward or rightward velocity. We thus created ‘directional CTCF tracks’ (CTCF ChIP-seq ‘painted’ with the directional sequence motif score, see Methods.)The directional CTCF tracks achieved the highest correlation and displayed the expected asymmetric effects in all cell types, with forward tracks reducing leftward velocity (*L*) and reverse tracks reducing rightward velocity (*R*) — an effect that can be observed when visually comparing the tracks as well (Fig. S4). Transcription-associated features such as H3K4me1, H3K4me3, and H3K36me3 also correlated with extrusion rates, but these effects were notably variable between cell types (Fig. 3B).

**Fig. 3:**
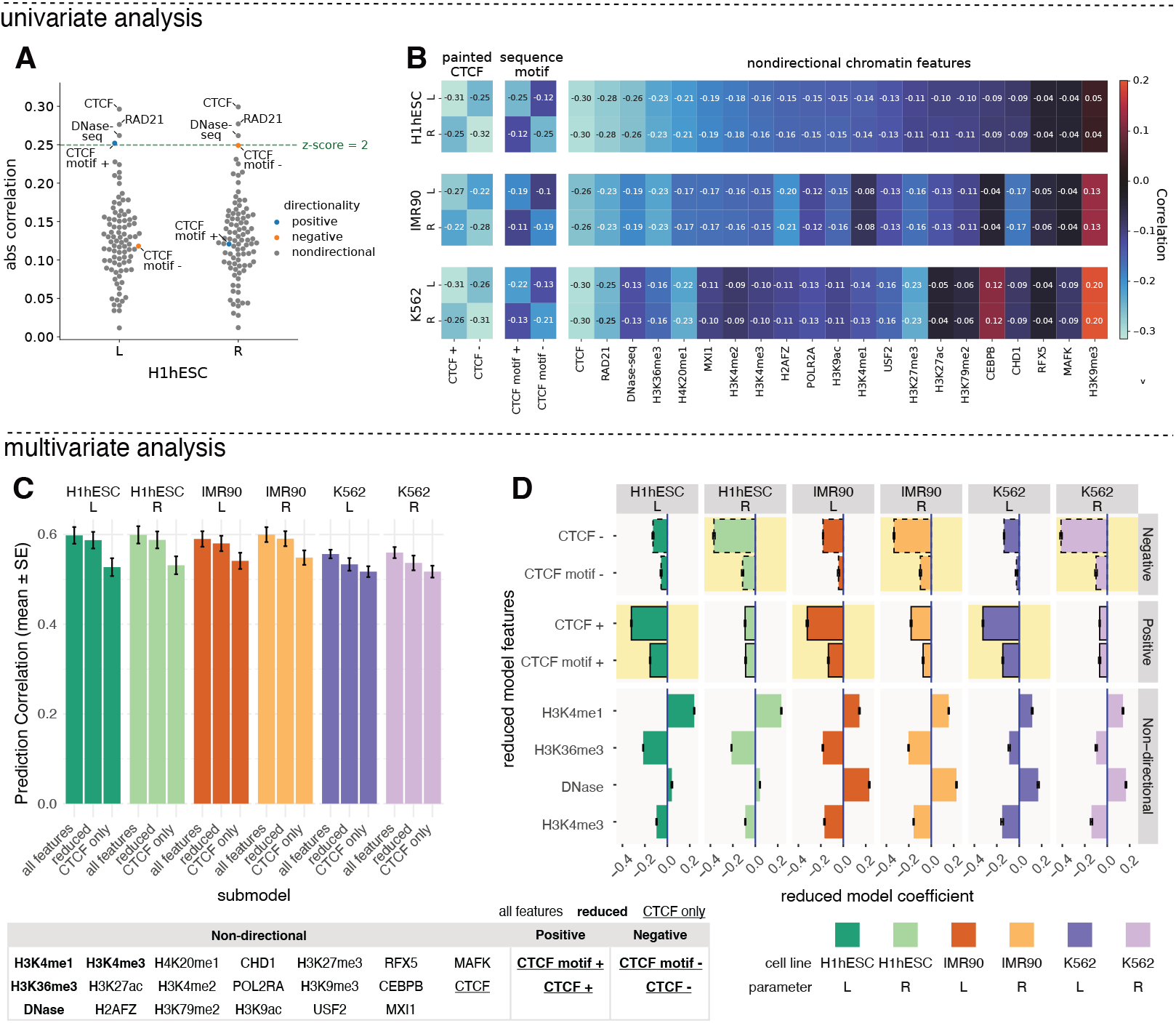
**A**: *Loop extrusion machinery tracks show strongest correlations with dLEM parameters*. Absolute correlations between fitted leftward (*L*) and rightward (*R*) velocity parameters and all available ENCODE ChIP-seq tracks for H1hESC (chromosomes 3-8 and 10-13, binned to 10kb resolution). Tracks exceeding z-score = 2 significance threshold are explicitly labeled, with loop extrusion machinery components (RAD21, CTCF) showing the highest correlations. CTCF motifs (+ and -) exhibit expected asymmetric correlations with *L* and *R* parameters, consistent with orientation-dependent blocking of cohesin translocation. **B**: *Univariate cross-cell-type analysis reveals consistent CTCF asymmetry and active transcription signals*. Heatmap showing univariate correlations between dLEM parameters and genomic tracks shared across three well-profiled cell lines (H1hESC, IMR90, K562). Loop extrusion machinery tracks (RAD21, CTCF) maintain consistently high correlations across all cell types. Directionally-signed CTCF (“painted” CTCF) tracks show consistent asymmetric patterns with *L* and *R* parameters across cell lines. While other tracks exhibit cell-type-specific variation, those with higher correlations are predominantly associated with active transcription, suggesting a broader role for transcriptional machinery in modulating cohesin dynamics. **C**: *Multivariate feature importance for predicting loop extrusion parameters*. Prediction performance across three cell lines and both parameters for models using all features, reduced features (CTCF plus top active transcription markers), and CTCF-only features (see feature table below). High performance of reduced model indicates significant contribution of active transcriptional machinery to loop extrusion dynamics. **D**: *Model coefficients of multivariate analysis reveal genomic determinants of loop extrusion*. Elastic net coefficients for the reduced model show expected CTCF directional asymmetry and consistent positive contributions from active transcription markers across all cell lines and parameters, revealing genome-wide relationship between transcriptional activity and loop extrusion.

To quantify contributions more systematically, we turned to multivariate modeling. Because the apparent influence of bound factors such as CTCF extends into neighboring bins, we first applied convolution kernels to genomic tracks, which improved correlations with velocity profiles and reproduced expected APA patterns (Fig. S6A–C). Having established this preprocessing step, we next used the resulting smoothed tracks as inputs to elastic net regression, evaluating three feature sets across all three cell types: all features, a reduced cross-cell-type set, and a CTCF-only baseline (Fig. 3C). As expected, CTCF-only models retained substantial predictive power in every cell type, but adding a small number of transcription-related features significantly improved performance. The reduced model, in particular, provided a highly simplified yet robust description that nearly matched the accuracy of the full model while remaining interpretable. Importantly, these models are fully predictive: in each cell type their outputs can be visualized as reconstructed contact maps (Fig. S6D), revealing how choices of feature set—CTCF-only versus reduced—translate into distinct 2D architectural predictions.

Examining coefficients from the reduced model (Fig. 3D) highlights two complementary contributions that were consistent across cell types. Directional CTCF features neatly separated into *L* and *R* impediments with uniformly negative coefficients, consistent with their barrier role. Transcription-related features contributed both positive and negative effects: H3K4me3 and H3K36me3 (promoters and gene bodies) slowed extrusion, consistent with polymerase interference, whereas H3K4me1 and open chromatin facilitated more rapid translocation. Notably, these latter effects are the opposite of their univariate correlations, highlighting how the multivariate framework disambiguate indirect associations and yields a simplified, mechanistically coherent model.

Together, these results show that dLEM enables compact, predictive models of extrusion dynamics in multiple cell types. Both the coefficients and the reconstructed contact maps provide insight, demonstrating that while CTCF remains the dominant regulator, transcriptional machinery makes reproducible contributions that become apparent when features are considered jointly.

### Interpretable models enable quantitative trans perturbation predictions

A central advantage of dLEM is that its parameters are physically interpretable, directly corresponding to extrusion dynamics such as detachment rate and velocity impediments. This structure enables not only descriptive and predictive modeling but also counterfactual reasoning about perturbations, a capability unavailable to black-box deep learning models. Global changes to loop extrusion can be represented by systematic shifts in parameter values, enabling quantitative trans-perturbation analysis. This means dLEM can predict how experimental perturbations will alter chromatin architecture and test mechanistic hypotheses quantitatively.

We illustrate this principle by examining two major perturbations: WAPL depletion and CTCF depletion (Fig. 4). For both, we leverage auxin-induced degradation datasets [26]. In this setting, dLEM can be applied in two complementary ways (Fig. 1F): first, to directly fit the perturbation data for descriptive analysis, and second, to predict perturbation effects by systematically manipulating parameters learned from the unperturbed condition.

**Fig. 4:**
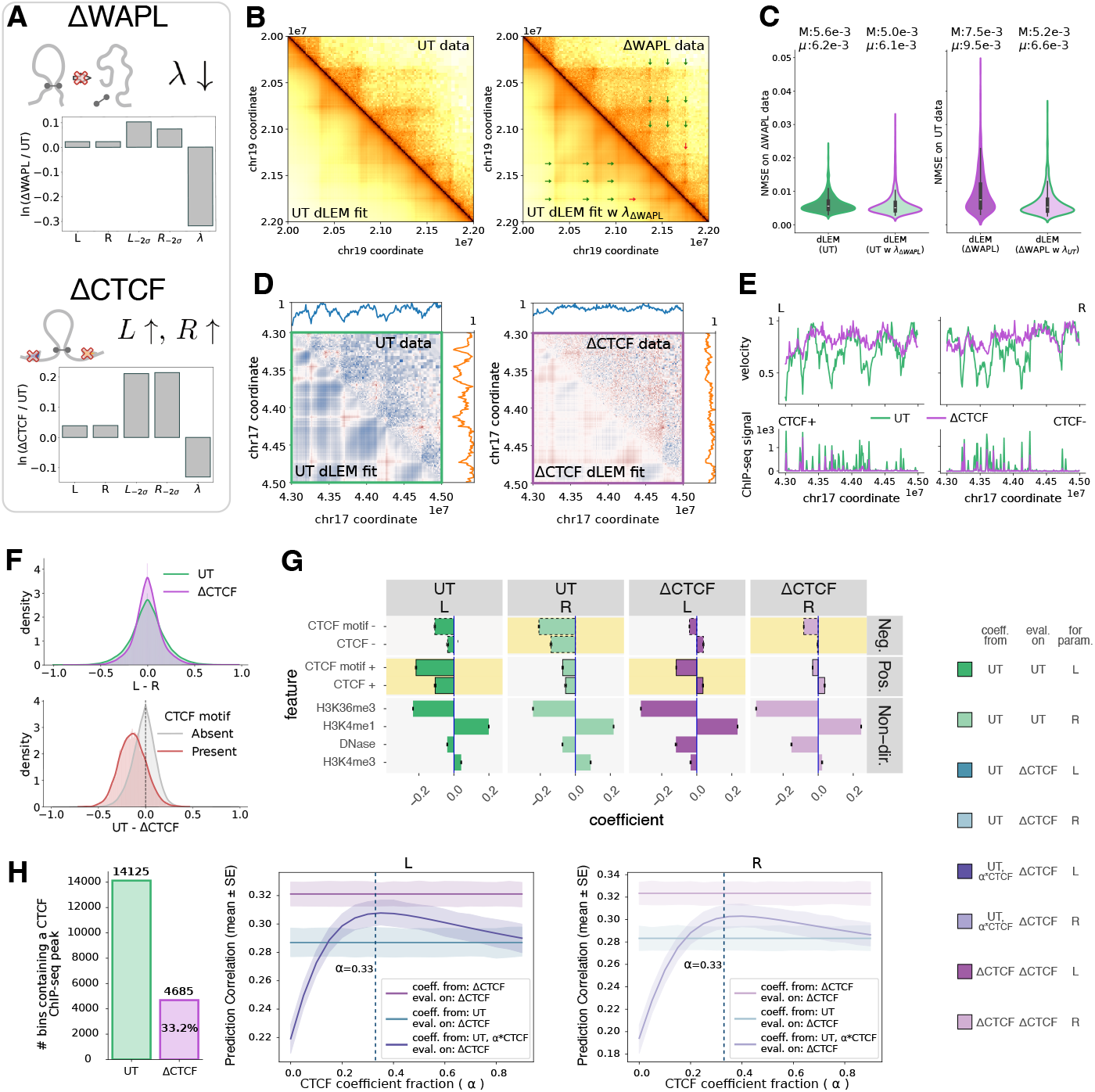
**A**: *dLEM predictions for loop extrusion perturbations*. Schematic showing expected effects of WAPL depletion (decreased detachment rate *λ*) and CTCF depletion (reduced velocity impediments *L, R*) on loop extrusion dynamics. Barplot shows fold changes in average dLEM parameters confirm predictions, with WAPL depletion primarily affecting *λ* and CTCF depletion primarily affecting *L* and *R* parameters (*L*_2*σ*_ and *R*_*−*2*σ*_ represent regions with the largest velocity dips). **B**: *WAPL depletion effects captured by detachment rate changes*. Contact map patch showing how switching *λ* values between untreated (UT) and ΔWAPL conditions affects loop formation predictions. Left: UT fit on UT data. Right: UT fit with *λ*_ΔWAPL_ compared to ΔWAPL data. Green arrows indicate additional loops captured by the modified model, red arrow indicates a loop not captured by the model. **C**: *Genome-wide validation of λ-mediated WAPL effects*. Normalized mean squared error (NMSE) across sliding window patches from 6 chromosomes. **D**: *dLEM captures CTCF depletion effects*. Representative genomic region showing experimental data (upper triangles) and dLEM fits (lower triangles) for untreated (UT) and ΔCTCF conditions. **E**: *CTCF depletion reduces velocity parameter impediments*. Velocity parameters *L* (top left) and *R* (top right) for the same region show reduced impediments (shallower dips) in ΔCTCF (purple) compared to UT (green). Bottom panels show directional ChIP-seq signal (CTCF+ left and CTCF- right) for the same genomic region. **F**: *CTCF depletion reduces asymmetry and affects motif-containing sites*. Top: Distribution of L-R differences shows reduced asymmetry in ΔCTCF compared to UT. Bottom: Parameter changes (UT - ΔCTCF) are concentrated at CTCF motif-containing sites, while motif-absent sites show minimal change. **G**: *Linear model coefficients reflect CTCF perturbation effects*. Elastic net coefficients for UT and ΔCTCF models show reduced CTCF directional asymmetry and compensatory increases in transcriptional machinery coefficients following CTCF depletion. **H**: *Interpretable prediction of perturbation effects*. Left: Barplot showing the number of genomic bins containing a CTCF ChIP-seq peak in UT and ΔCTCF conditions, with 33.2% of peaks remaining after auxin treatment. Center and right: Correlation between predicted and fitted dLEM parameters (*L* and *R*) for the ΔCTCF condition across three model scenarios: UT model trained and evaluated on ΔCTCF data (green), ΔCTCF model trained and evaluated on ΔCTCF data (purple), and UT model with systematically reduced CTCF coefficients (fraction *α* on x-axis) evaluated on ΔCTCF data (blue). *α* = 0.33, corresponds to the experimentally observed fraction of remaining CTCF peaks.

Given the known function of WAPL as a cohesin unloader, we can predict a specific effect of WAPL depletion in dLEM: a decrease in the detachment rate *λ* (Fig. 4A), while leaving other parameters (velocity profiles) unchanged. Because the influence of *λ* is difficult to visualize on the observed-over-expected scale, we present results on the absolute scale. The observed features—extended loop-associated stripes, a denser grid of dots, and increased distal contact mass—are consistent with reduced cohesin release and underscore WAPL’s role in regulating residence time (Fig. 4B) [26, 27]. Importantly, WAPL is a highly constrained gene (gnomAD pLI = 1) [28], suggesting dosage sensitivity with potentially large phenotypic consequences downstream of subtle architectural effects. This indicates a strong selection against loss-of-function variants and suggests dosage sensitivity with potentially large phenotypic consequences. Even modest changes in cohesin residence time could have downstream effects on gene regulation and development, making quantitative prediction of WAPL perturbation effects clinically relevant.

To test whether WAPL depletion indeed affects primarily the detachment rate, we performed the following experiment: we separately fit models to UT and ΔWAPL data, then replaced only the detachment rate from one condition into the other. Predicting ΔWAPL from UT via a single scalar parameter change, we successfully recapitulated the majority of newly emerged dots (Fig. 4B). For completeness, we also reversed the process, applying *λ*_UT_ to the ΔWAPL model, with both directions demonstrated on additional patches in Fig. S7. Genome-wide evaluation confirmed improved model fits from single parameter swap (Fig. 4C), as measured by normalized mean squared error (NMSE), a more appropriate metric on the absolute scale since correlations are dominated by diagonal decay and remain uniformly high.

In contrast to ΔWAPL, the ΔCTCF condition is expected to affect the extrusion velocity parameters *L* and *R*. Specifically, dLEM predicts that loss of CTCF impediments reduces *L* and *R* asymmetry and weakens boundary insulation, which is precisely what we observe in the fitted parameters and contact maps.(Fig. 4D–F).

The descriptive elastic net analysis introduced in Fig. 3 provides complementary support: when refit under ΔCTCF, CTCF-associated coefficients diminish and features related to transcriptional machinery become more prominent (Fig. 4G). Together, these results illustrate how the descriptive models from Fig. 2–3 can be leveraged predictively: perturbation effects emerge directly from shifts in interpretable parameters.

Most importantly, dLEM’s interpretable structure enables quantitative testing of mechanistic hypotheses about perturbations. While auxin-induced degradation is typically assumed to achieve complete protein depletion, CTCF ChIP-seq data reveals that 33% of significant peaks persist in the ΔCTCF condition at 10kb resolution, and fitted dLEM velocity parameters show correspondingly weakened but persistent impediments at these sites (Fig. 4E). A key question is whether this is due to residual CTCF remains functionally capable of blocking cohesin translocation.

To test this, we trained elastic net models using only UT-fitted dLEM parameters and corresponding UT CTCF and other epigenetic tracks, then systematically scaled the CTCF-related coefficients by a factor *α* (0–1) evaluating predictive performance on ΔCTCF-fitted dLEM parameters (Fig. 4H). Setting *α* = 0 substantially degraded predictions below the model trained on UT data, indicating that some CTCF contribution is still required. Strikingly, maximal performance was achieved at *α ≈* 0.33, closely matching the experimentally observed fraction of retained CTCF peaks in the ΔCTCF condition—data that were not used in model training. Larger *α* values reduced performance, indicating that residual CTCF operates at reduced but non-zero functional capacity. This quantitative correspondence shows that residual CTCF retains functional barrier activity. More broadly, this analysis exemplifies how dLEM’s interpretable parameters enable principled perturbation testing unavailable to black-box models. By systematically varying a single interpretable parameter (*α*), we quantitatively determined the functional capacity of residual CTCF. This capacity for mechanistic hypothesis testing distinguishes dLEM from data-driven approaches where model parameters cannot be meaningfully perturbed to generate testable biological predictions.

### Deep dLEM provides a predictive framework for 3D genome architecture

A central design goal for dLEM was not only to provide a descriptive tool but also a biophysical layer for prediction. Linear models relating *L* and *R* parameters to genomic tracks hint at this potential, but they are limited in power and depend heavily on CTCF ChIP–seq, which is not always available. To overcome these constraints, we developed deep dLEM, a deep learning architecture that infers extrusion parameters from local sequence and chromatin context and then passes them through a fixed mechanistic head to generate 2D contact maps. This design anchors the model in physics while allowing upstream neural components to flexibly discover informative features.

By embedding the mechanistic dLEM layer as a fixed component, deep dLEM achieves a two-fold advantage: unlike black-box models that must learn the 1D-to-2D mapping implicitly through millions of parameters, deep dLEM hard-codes this transformation through the biophysical model (dLEM), simultaneously maintaining interpretability and dramatically reducing model complexity by eliminating the need for parameters devoted solely to learning this mapping.

Our guiding principle was to enable context-specific prediction from minimal and widely available data. For broad experimental deployability, deep dLEM requires only DNA sequence and open chromatin accessibility (e.g., ATAC-seq or DNaseseq). Because CTCF binding nearly always coincides with open chromatin, combining motif orientation from sequence with accessibility signals provides a reliable proxy for functional occupancy. This makes the model immediately applicable even when comprehensive ChIP-seq data are unavailable – a common limitation in clinical samples, rare cell types, developmental time courses, or resource-constrained settings. This balance of minimal data requirements and high predictive accuracy makes deep dLEM broadly deployable across diverse experimental contexts.

Deep dLEM takes two types of input, a non-contextual sequence representation and cell type–specific accessibility profiles at 8kb resolution, which are processed by a compact neural network that outputs *L* and *R* extrusion rates. These rates are then transformed into 2D observed-over-expected contact maps by the fixed dLEM head, and the entire model is trained end-to-end with mean squared error loss (Fig. 5A).

**Fig. 5:**
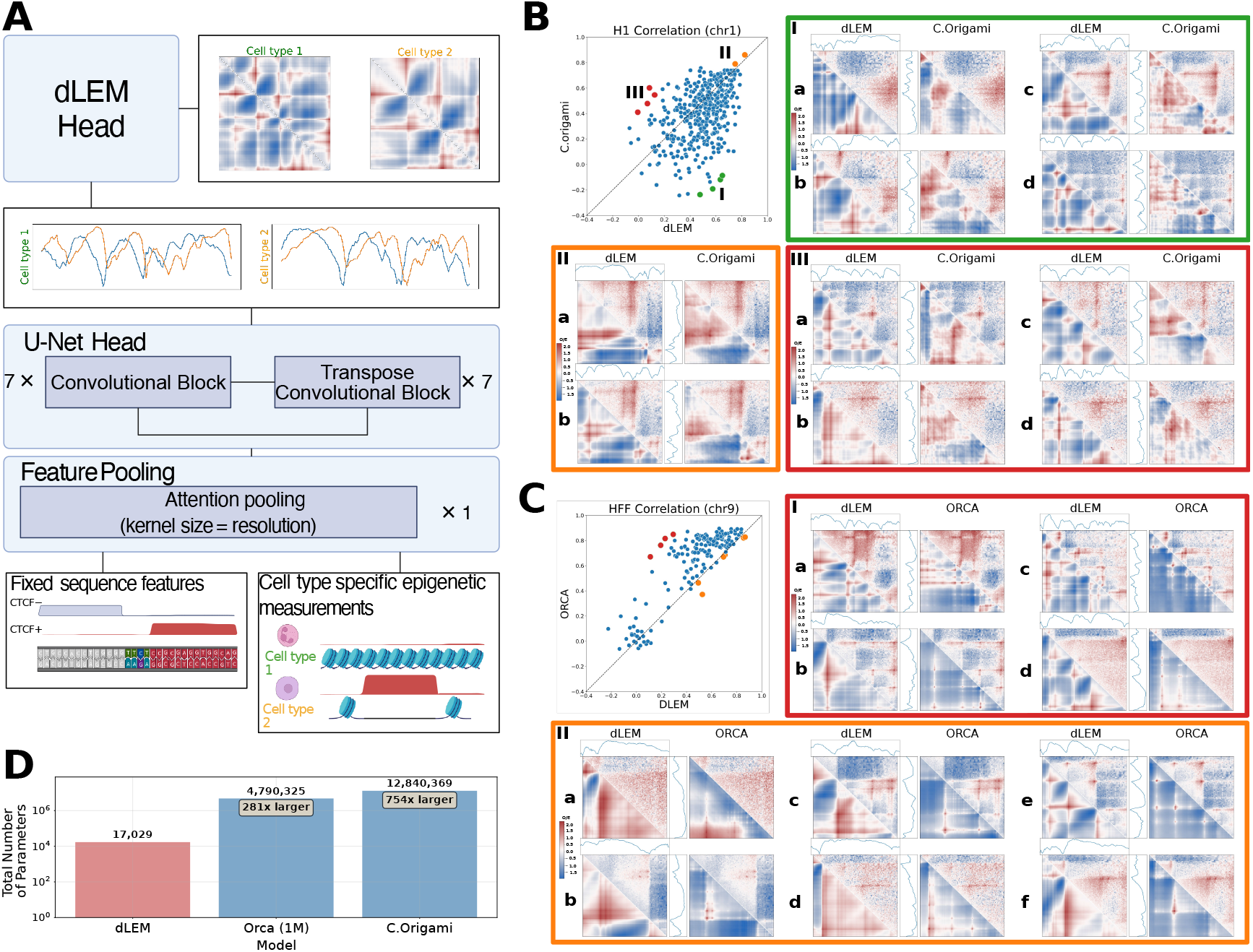
**A**: *Architecture of the deep dLEM model*. Deep dLEM takes two types of input: a non-contextual sequence representation and cell type–specific epigenetic measurements that provide contextual information. These 1D inputs are processed by a small neural network that outputs two rate parameters, which are then passed through a fixed dLEM head to generate 2D observed-over-expected contact maps. The model is trained end-to-end using mean squared error (MSE) loss against experimental data. **B**: *Comparison of dLEM and C*.*Origami*. We compare dLEM with C.Origami, another explicitly contextualized model. Both models are trained on HFF data and used to predict H1 data at matched loci. Performance is assessed by Pearson correlation between observed and expected values. Overall, dLEM achieves slightly higher correlations on average. Example patches from C.Origami *>* dLEM (red), C.Origami *<* dLEM(green), and C.Origami ≈ dLEM (orange) performance regimes are shown. **C**: *Comparison of dLEM and Orca*. dLEM is evaluated against Orca, a high-performing model trained on the same Micro-C input data. Both models are assessed on identical validation patches. As expected, Orca outperforms dLEM overall; however, patch-level performance is strongly correlated across models, indicating that prediction difficulty is shared. Example patches from different performance regimes are highlighted as in Panel B. **D**: *Model complexity*. Parameter counts for the three models demonstrate that deep dLEM is orders of magnitude smaller than both C.Origami and Orca, highlighting its lightweight architecture.

We compare deep dLEM against C.Origami [23], which also combines sequence with cell-type context but explicitly requires measured CTCF ChIP-seq as input. Trained on HFF and evaluated on H1, both models showed broadly similar correlations, with most loci clustering near the diagonal (Fig. 5B). Yet more loci fell in the region where deep dLEM outperformed, and—because its 1D-to-2D mapping is hard-coded through the dLEM layer—deep dLEM required orders of magnitude fewer parameters to achieve this (Fig. 5 D).

For context within the broader landscape of 3D structure prediction, we also compared deep dLEM to Orca [22]. Unlike C.Origami, Orca is unconditional and cannot be used to predict across cell types but benefits from being trained on the same Micro-C data used for deep dLEM, and it is highly performant. On loci present in the test set for both models, Orca unsurprisingly outperformed deep dLEM in same-cell-type, different-locus predictions, though performance remained strongly correlated across loci (Fig. 5C).

Inspection of patches with the largest discrepancies between the two models (red border) revealed distinct modes of performance. In two regions (a, b), deep dLEM correctly identified loop anchors but underestimated insulation strength, while in another (c) it missed a subset of anchors. In a fourth case (d), both models incorrectly predicted anchors in an otherwise featureless TAD, though Orca produced a sharper boundary. By contrast, in regions of comparable performance (orange border), both models generally agreed on anchor placement—even reproducing the same errors—yet often differed in how they scaled intra- versus inter-TAD interactions. Additional examples covering the full range of model behaviors are provided in Fig. S8 and Fig. S9.

Despite these differences, deep dLEM achieves comparable predictive behavior using 700× fewer parameters than Orca (Fig. 5 D). Combined with its cross-cell-type performance relative to C.Origami, this level of model compression suggests that much of the representational burden in large chromatin folding models is devoted to implicitly learning loop extrusion dynamics—structure that deep dLEM encodes explicitly through its biophysical layer.

Taken together, these benchmarks establish deep dLEM as a lightweight yet powerful predictive framework. By embedding extrusion dynamics as a fixed biophysical layer, deep dLEM unites the parsimony and interpretability of a mechanistic model with the flexibility of modern deep learning. This design achieves competitive accuracy with two to three orders of magnitude fewer parameters than existing approaches. More broadly, deep dLEM points to a new paradigm: using biophysics as an architectural prior to build compact, transparent, and generalizable models of genome folding and genomic processes more broadly.

## Discussion

The differentiable loop extrusion model (dLEM) provides a genome-wide, mechanistically interpretable framework for connecting one-dimensional genomic features to three-dimensional chromosome architecture. By reducing complex contact maps to simple extrusion parameters, dLEM isolates the contribution of cohesin-mediated loop extrusion, enabling quantitative predictions of architectural changes under perturbation and direct comparison across cell types. Deep dLEM further extends this framework, showing that extrusion parameters can be inferred directly from sequence and chromatin features with high accuracy and efficiency. Together, these results establish loop extrusion velocity as a tractable latent space for predictive modeling and provide a foundation for systematic dissection of the mechanisms that shape genome organization.

While dLEM focuses on a single physical mechanism, it offers a blueprint for extending mechanistic modeling of genome folding. Incorporating additional processes such as compartmental organization, explicit interactions with transcriptional machinery, and spatially regulated cohesin turnover will further enrich the framework. Looking forward, the ability to embed extrusion within a broader multi-process model of genome architecture holds promise for unifying distinct scales of chromatin folding. In this way, dLEM not only clarifies the role of loop extrusion but also opens the door to a new generation of interpretable, predictive models of 3D genome organization.

## Methods

For opening and processing the Hi-C/Micro-C contact maps, we use cooltools [29]. For dLEM fitting and deep dLEM development we use PyTorch [30]. All analyses and data generation in this work were performed using dLEM, which is available as a Python package for easy installation and use in other projects. The software for directly fitting dLEM to contact map data is available at https://github.com/chikinalab/dLEM and deep dLEM model is available at https://github.com/chikinalab/dLEMpytorch.

### Fitting the dLEM “direct fit” model

#### Fitting contact-decay model

To obtain the contact-decay data for each chromosome, we collected the average value of each diagonal of the balanced Hi-C/Micro-C matrix up to 3.75Mb (giving a vector of 375 diagonals for 10kb resolution). To fit the contact-decay model (Fig. 1E) we omitted the first 7 diagonals, as they exhibit high contact frequency, part of which may come from technical biases or local chromatin interactions which cannot be captured by our model. To obtain the vector of the genomic distances between loci, *x*, we assumed the value of the center of the diagonal vector bins. To find model parameters *a, b, c*, and the ratio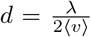, the model was fitted to the data using L-BFGS (Limited-memory Broyden–Fletcher–Goldfarb–Shanno) algorithm, MSE (Mean Squared Error) loss between the log data and log model prediction, 200 epochs, and initial parameters *a* = 0.05, *b* = 1.1, *c* = 0.1, and *d* = 0.01.

#### Generating a contact map using dLEM

Consider a contact map patch spanning loci 0, …, *j*, …, *i*, …, *N*. To generate the contact frequency at pixel *C*^*i,j*^, we require the velocity parameters at current and preceding-neighbor positions (*L*^*j*^, *L*^*j*+1^, *R*^*j*^, *R*^*i−*1^), the detachment rate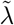, and the contact frequencies at adjacent pixels from the previous diagonal (*C*^*i,j*+1^ and *C*^*i−*1,*j*^). All indices *i, j, i −* 1, and *j* + 1 remain within the patch boundaries [0, *N*]. The computational structure follows the diagonal organization of the contact map, which reflects the progressive extension of cohesin loops during extrusion. Since pixels *C*^*i,j*+1^ and *C*^*i−*1,*j*^ are located on the preceding diagonal, all pixels on any given diagonal *d* can be computed simultaneously and depend only on the values from diagonal *d−*1. This diagonal-by-diagonal generation requires three inputs: the velocity vectors *L* and *R*, the scalar detachment rate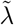, and the contact frequencies from the previous diagonal.

This formulation enables flexible contact map reconstruction. One can generate a complete contact map by starting from a uniform main diagonal and proceeding sequentially, or alternatively, generate any specific diagonal provided the preceding diagonal values are available.

#### Pre-processing contact map patch

Contact map data pre-processing serves two purposes: providing input diagonals for dLEM contact map generation and establishing target data for model fitting. Several challenges must be addressed before fitting dLEM to experimental data. First, contact map matrices contain substantial noise, particularly at higher resolutions, requiring careful handling of zero values that become problematic under logarithmic transformation. Second, since the core dLEM model describes loop extrusion dynamics without underlying polymer physics, we must isolate the extrusion signal from the broader contact frequency patterns to the extent possible.

Our preprocessing pipeline uses cooltools to: (1) balance the contact matrix to correct for experimental biases, (2) interpolate missing values, (3) coarsegrain the data using default parameters to reduce noise for log-space operations, (4) apply logarithmic transformation to improve signal resolution, and (5) normalize to obtain observed-over-expected contact frequencies by subtracting the expected background (diagonal averages) in log space. For direct model fitting, we transform the log-space observed-over-expected values back to linear scale through exponentiation to serve as input diagonals.

#### Fitting dLEM to one contact map patch

The optimization process to find vectors *L* and *R* takes as input the preprocessed observed-over-expected contact frequencies (in linear scale), initial velocity vectors *L*_0_ and *R*_0_ (initialized as uniform vectors of ones), and the detachment rate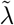. The detachment rate is either fitted from contact-decay analysis or approximated as 2.5 *×* 10^*−*6^ *h* (giving 0.025 for resolution *h* = 10*kb* and 0.005 for *h* = 2*kb*), a value which matches general range of values observed in contact-decay analysis.

Loop extrusion features are most prominent at distances between 100 kb and 1 Mb. We exploited dLEM’s flexibility to make predictions from any diagonal to restrict training to this range. For each input diagonal between *d*_start_ and *d*_end_, we (1) used the observed-over-expected values as initialization, (2) generated the next *k* diagonals from current *L* and *R* parameters, (3) computed MSE loss between log-scale predictions and targets for each diagonal, and (4) summed these losses across diagonals and starting positions. We used diagonals from 50 kb to 1.2 Mb with *k* = 10 for *h* = 10 kb, and from 25 to 600 with *k* = 50 for *h* = 2 kb (Fig. 2C).

Optimization proceeds for 100 epochs using the Adam optimizer, with *L* and *R* values clamped between 10^*−*9^ and 1 to maintain physical constraints. While MSE guides parameter updates, we independently track correlation and select the parameter set achieving the highest correlation rather than lowest MSE, as correlation better captures the structural patterns relevant to loop extrusion dynamics. Direct-fit dLEM provides a descriptive factorization of the contact map, similar to PCA, and therefore does not involve a cross-validation step.

#### Reconstructing full contact frequencies

The dLEM model captures only loop extrusion dynamics, while experimental contact maps reflect contacts from multiple sources. To reconstruct the full contact frequency scale, we combine the dLEM output with the distance-decay model fitted earlier. We assume that at the length scales of interest, contacts arise primarily from two independent processes: passive polymer diffusion (following a power-law decay) and active loop extrusion.

To generate the scaled contact map (Fig. 2B, we: (1) create a distance-decay matrix based on the power-law component with exponent *b*, normalized so the main diagonal equals one, (2) generate the loop extrusion component using dLEM with the main diagonal also set to one, (3) combine these components as *c* · dLEM + power-law, where *c* is the fitted loop extrusion contribution coefficient from the distance-decay analysis, and (4) scale the combined result to match experimental data magnitude using the fitted scaling parameter *a*. This approach separates the relative contributions of different physical processes while maintaining quantitative agreement with observed contact frequencies.

#### Fitting dLEM to a whole chromosome

To estimate dLEM parameters across an entire chromosome, we employ a sliding window approach with overlapping patches. This strategy stabilizes parameter estimation by allowing each genomic region to contribute to multiple patch fits, enabling robust loop extrusion dynamics to emerge while averaging out patch-specific noise variations. However, parameter estimation quality varies by position within each patch: parameters at patch centers contribute to a larger portion of the contact map and thus receive more robust estimates than those at patch edges, where parameters influence only a limited number of contact predictions.

Additionally, chromosomal regions with sparse contact map coverage pose estimation challenges, as patches spanning low-coverage areas provide less reliable parameter constraints. To address these issues, we compute final *L* and *R* values using a weighted average across all overlapping patches. The weight for each parameter estimate combines two factors: (1) patch coverage, calculated as the percentage of non-NaN values along the main diagonal within the patch, and (2) positional centrality, reflecting how close the parameter’s genomic coordinate is to the center of the patch. This weighting scheme prioritizes high-quality estimates from well-covered regions while maintaining parameter continuity across the chromosome.

To generate a chromosome-wide fits used in Fig. 2, Fig. 3, and Fig. 4, we used patch size 2Mb (200 *×* 200 bins for *h* = 10*kb*) and sliding window 500kb (50 bins for *h* = 10*kb*), giving 4 values for *L* and *R* at each position, which were then averaged using before-mentioned weights to arrive to the final estimates for *L* and *R*.

#### Fitted cell lines

For figures 2-4, we analyzed Hi-C and Micro-C data (.mcool files from 4D Nucleome) from four human cell lines (Table 1). Our analysis focused on chromosomes 3-8 and 10-13, excluding chromosome 9 due to extensive regions lacking coverage that could potentially bias the results of the downstream analysis.

**Table 1:**
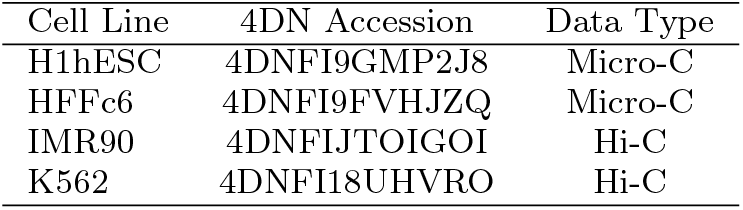
human Hi-C and Micro-C datasets used in this study.

#### Generating chromosome-wide patch correlations between fit and data

To evaluate dLEM performance across diverse genomic contexts for the violin plots in Figure 2E, we generated correlations using a sliding window approach with 200kb step size (20 bins) and 2Mb patch size (200 bins) across chromosomes 3-8 and 10-13. The detachment rate 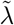 was fitted individually for each chromosome using the contact-decay model to account for chromosome-specific characteristics. For each patch, we computed Pearson correlations between the flattened observed-over-expected experimental data and the corresponding flattened dLEM predictions, providing a comprehensive assessment of model performance across varied chromatin architectures and genomic contexts.

### ENCODE track processing

To compare genomic tracks with dLEM parameters, we processed publicly available genomic data to match the resolution and length of the fitted velocity parameters. We downloaded ChIP-seq, DNase-seq, and RNA-seq data in bigWig format from the ENCODE database for three well-profiled cell lines (H1hESC, IMR90, and K562). All tracks were binned to 10kb resolution using both mean and maximum summary statistics to capture different aspects of the signal distribution within each bin.

To generate directional CTCF tracks (referred to as CTCF+ and CTCF- or “painted CTCF”), we multiplied the binned CTCF ChIP-seq signal with the corresponding binned CTCF motif orientation tracks (mean or max summary of the bin), creating strand-specific CTCF binding profiles. Following binning, all tracks were convolved with optimized kernels specific to *L* and *R* parameters, as described in the linear modeling section.

For experiments with multiple replicates or summary statistics, we selected the track yielding the highest correlation with dLEM parameters. For non-directional features, we chose the track with the highest mean absolute correlation across both *L* and *R* parameters. For directional CTCF features, we selected tracks showing the strongest correlation in the expected orientation (positive correlation with *L* parameters and negative correlation with *R* parameters), consistent with the known asymmetric blocking effects of convergent CTCF binding sites. The final selected tracks and their ENCODE accession numbers are listed in Tables 2, 3, and 4.

**Table 2:** Selected ENCODE tracks for H1hESC cell line.

| Track | ENCODE Accession | Summary |
| --- | --- | --- |
| H3K9me3 | ENCFF441SLB | mean |
| H3K36me3 | ENCFF985CVI | max |
| H3K27ac | ENCFF771GNB | max |
| H2AFZ | ENCFF519VGB | max |
| H4K20me1 | ENCFF674EST | max |
| USF2 | ENCFF857KTJ | mean |
| RFX5 | ENCFF495SBQ | mean |
| CEBPB | ENCFF258EZO | mean |
| H3K4me3 | ENCFF726NIH | max |
| H3K4me2 | ENCFF556VEB | max |
| MXI1 | ENCFF126KRU | mean |
| H3K4me1 | ENCFF753ZAB | max |
| DNase-seq | ENCFF232GUZ | max |
| H3K79me2 | ENCFF868TLW | max |
| CHD1 | ENCFF078DNM | max |
| POLR2AphosphoS5 | ENCFF491XDR | max |
| H3K27me3 | ENCFF395GVR | max |
| MAFK | ENCFF959KBD | mean |
| H3K9ac | ENCFF354HBK | max |
| RAD21 | ENCFF837RSL | max |
| CTCF | ENCFF269OPL | max |
| CTCF motif + | - | max |
| CTCF motif - | - | max |
| painted CTCF + | ENCFF269OPL | max ChIP-Seq mean motif+ |
| painted CTCF - | ENCFF269OPL | max ChIP-Seq mean motif- |

**Table 3:** Selected ENCODE tracks for IMR90 cell line.

| Track | ENCODE Accession | Summary |
| --- | --- | --- |
| H3K9me3 | ENCFF021HVH | mean |
| H3K36me3 | ENCFF968ONU | max |
| H3K27ac | ENCFF694JJX | max |
| H2AFZ | ENCFF905MQV | max |
| H4K20me1 | ENCFF466TGZ | max |
| USF2 | ENCFF186TGJ | mean |
| RFX5 | ENCFF039PUG | mean |
| CEBPB | ENCFF342JPR | mean |
| H3K4me3 | ENCFF433XYT | max |
| H3K4me2 | ENCFF195RXD | max |
| MXI1 | ENCFF097YGG | mean |
| H3K4me1 | ENCFF806SLL | max |
| DNase-seq | ENCFF323RPM | max |
| H3K79me2 | ENCFF623CHU | max |
| CHD1 | ENCFF793YSZ | max |
| POLR2A | ENCFF616IEH | max |
| H3K27me3 | ENCFF625XOO | max |
| MAFK | ENCFF551ANQ | mean |
| H3K9ac | ENCFF431URC | max |
| RAD21 | ENCFF696GJW | max |
| CTCF | ENCFF863CKA | max |
| CTCF motif + | - | max |
| CTCF motif - | - | max |
| painted CTCF + | ENCFF863CKA | max ChIP-Seq mean motif+ |
| painted CTCF - | ENCFF863CKA | max ChIP-Seq mean motif- |

**Table 4:** Selected ENCODE tracks for K562 cell line.

| Track | ENCODE Accession | Summary |
| --- | --- | --- |
| H3K9me3 | ENCFF559MMQ | mean |
| H3K36me3 | ENCFF811PDH | max |
| H3K27ac | ENCFF289FLK | max |
| H2AFZ | ENCFF903BRE | max |
| H4K20me1 | ENCFF990ELV | max |
| USF2 | ENCFF235XFA | mean |
| RFX5 | ENCFF028COD | mean |
| CEBPB | ENCFF978UZJ | mean |
| H3K4me3 | ENCFF307STR | max |
| H3K4me2 | ENCFF623KCP | max |
| MXI1 | ENCFF845HTM | mean |
| H3K4me1 | ENCFF948OCT | max |
| DNase-seq | ENCFF972GVB | max |
| H3K79me2 | ENCFF839BCY | max |
| CHD1 | ENCFF184FBU | max |
| POLR2AphosphoS2 | ENCFF060UOO | max |
| H3K27me3 | ENCFF521GNI | max |
| MAFK | ENCFF556ZFU | mean |
| H3K9ac | ENCFF806RDN | max |
| RAD21 | ENCFF666JJM | max |
| CTCF | ENCFF525FLS | max |
| CTCF motif + | - | max |
| CTCF motif - | - | max |
| painted CTCF + | ENCFF525FLS | max ChIP-Seq mean motif+ |
| painted CTCF - | ENCFF525FLS | max ChIP-Seq mean motif- |

#### Track convolution with local chromatin architecture kernel

To account for the local effects of chromatin features on loop extrusion parameters, we learned feature-specific convolution kernels. The goal was to find kernels that maximize the correlation between convolved genomic tracks and dLEM parameters while preserving the directionality of the original correlation. For each genomic feature, we optimized a convolution kernel *k* by minimizing the loss function:

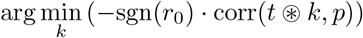

where *k* is the kernel, sgn(*r*_0_) is the sign of the original correlation between the non-convolved track and parameter, *t* is the genomic track, *p* is the dLEM parameter (*L* or *R*), and ⊛ denotes convolution. This formulation ensures that kernel optimization enhances correlation strength while maintaining the original positive or negative relationship between the feature and parameter. We enforced non-negativity constraints on all kernel weights to ensure biological interpretability. Kernels had a fixed width of 25 bins and were initialized with a Gaussian distribution centered at the kernel midpoint with variance 1, plus small random noise. Optimization proceeded for 250 epochs using Adam optimizer.

#### Comparison of kernels with aggregated peaks

To validate that learned convolution kernels capture biologically meaningful local chromatin architecture effects, we compared kernel predictions with experimental aggregate peak analysis (APA) patterns.

For each genomic track, we identified peak locations using a z-score threshold of 2.5 and generated paired coordinates using `bioframe.pair by distance()` with minimum separation of 500kb and maximum separation of 1.2Mb. These coordinates were used to create pileups with `cooltools.api.snipping.pileup()` using an 800kb flank around each peak pair. Pileups were generated on the original contact frequency scale rather than observed-over-expected. The large flank size ensured sufficient baseline regions for robust marginal calculations. Observed-over-expected normalization was applied post-pileup through diagonal normalization of the aggregated contact matrix.

APA marginals were computed by summing the APA plot along the X and Y dimensions. To generate convolved peak comparisons, we created average peak shapes by averaging the top peaks for each track, then convolved these average peaks with the optimized kernels. The “uncoupled APA” reference was generated as the outer product of the X and Y APA marginals, representing the expected pattern from independent marginal distributions. Finally, “kernel predictions” were computed as the outer product of the *L* and *R* convolved peaks.

### ENCODE track analysis

To investigate whether dLEM parameters can be predicted from ENCODE tracks, we constructed a series of linear models using the glmnet package in R. For each model, the “left” convolved tracks were used to predict the left dLEM parameter, while the “right” convolved tracks were used for the right dLEM parameter. Three different sets of features were used for training: all features, reduced (DNase, H3K36me3, H3K4me1, H3K4me3, CTCF painted +, CTCF painted -, CTCF motif +, CTCF motif -) and CTCF related (CTCF, CTCF painted +, CTCF painted -, CTCF motif +, CTCF motif -). Prior to training, both the input tracks and the target parameters were scaled. Model training was performed using 5-fold cross-validation to assess the stability of the estimated coefficients and the robustness of prediction performance. We applied elastic net regularization with a fixed mixing parameter of *α* = 0.9, while allowing the regularization strength *λ* to be optimized during training. Model quality in each fold was evaluated by computing the correlation between the predicted values and the corresponding dLEM parameters.

### Trans-perturbation experiments

For analysis of trans-perturbation effects on 3D chromatin structure, we used publicly available Micro-C and ChIP-seq data from Hsieh *et al*. [26] available under Gene Expression Omnibus (GEO) accession GSE178982 (Table 5).

**Table 5:**
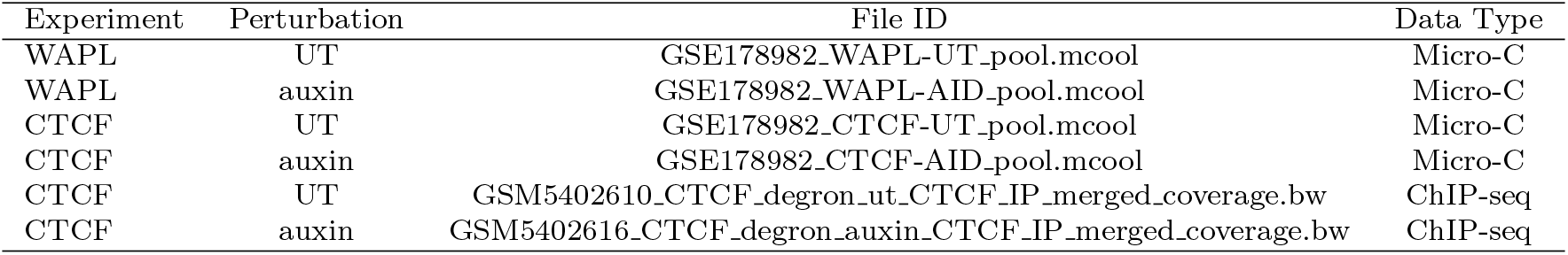
Trans-perturbation datasets used in this study.

We generated direct dLEM fits for both untreated (UT) and auxin-treated conditions across multiple chromosomes using approach described in Paragraph 5. We fitted the detachment rate parameter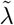 individually for each condition and chromosome using the contact-decay model, then applied the direct fitting procedure described above to estimate velocity parameters *L* and *R*. For WAPL perturbation experiments, we analyzed chromosomes 14-19, while for CTCF perturbation experiments, we analyzed chromosomes 8-19.

To study the fold change of average parameters between UT and auxin-treated conditions for both WAPL and CTCF perturbations, we first computed the average parameters ⟨*L*⟩, ⟨*R*⟩ and average *λ* across chromosomes, where *λ*^chrom^ for each chro-mosome was computed as 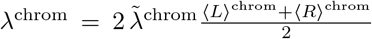. The fold change was computed as the ratio of average parameters in treated over untreated conditions.

However, since velocity impediments (dips in *L* and *R* parameters) are sparse by construction, genome-wide averages may not capture the full extent of perturbation-induced changes. To address this limitation, we additionally analyzed parameters with the most pronounced velocity impediments, defined as positions with z-score ≤ *−*2 relative to the chromosome-wide parameter distribution. This threshold-based analysis allows us to assess how perturbations specifically affect regions with strong loop extrusion impediments, which are most likely to reflect biologically relevant cohesin dynamics.

### WAPL trans-perturbation

To test the hypothesis that WAPL perturbation effects can be largely explained by changes in the detachment rate *λ*, we performed a parameter transfer experiment. We fitted dLEM parameters separately to UT and ΔWAPL conditions, then systematically exchanged the *λ* values between conditions while keeping all other parameters unchanged.

The parameter transfer required converting between the exponential term exponent *d* used in the distance-decay model and 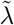 used in dLEM fitting, both of which are functions of *λ*, but *d* depends on average velocity while 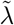 depends on maximum veloc-ity. For each condition, we computed λ = 2*d* ⟨*υ*⟩, where 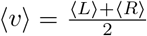 represents the average velocity for that condition. We then substituted *λ*_ΔWAPL_ into the UT model (creating “UT with *λ*_ΔWAPL_”) and *λ*_UT_ into the ΔWAPL model (creating “ΔWAPL with *λ*_UT_”). All other distance-decay model parameters (power-law exponent, scaling factors) remained specific to each condition.

To evaluate whether this single parameter change improves model accuracy, we compared the transferred models against their respective target data using normalized mean squared error (NMSE) on log-scale contact frequencies. We assessed performance across multiple genomic regions patches, using sliding window with 200Mb shift across chromosomes 14-19 to ensure robust statistical evaluation across chromosomes.

### CTCF trans-perturbation

For CTCF perturbation analysis, we generated direct dLEM fits for both untreated (UT) and auxin-treated conditions as described above. To analyze the relationship between CTCF motif presence and parameter changes induced by CTCF depletion, we computed residuals (UT - ΔCTCF) for both L and R parameters at each genomic position. We used positive-strand motifs used for L parameter analysis and negative-strand motifs used for R parameter analysis, reflecting the directional nature of CTCF blocking. We converted motif scores to z-scores and classified genomic bins as “motif present” (z-score ≥ 2) or “motif absent” (z-score *<* 2) to create binary categories for analysis.

#### Linear model analysis

For the mouse experiment, the maximum value of each track (Table 6) was computed, and the best replicate was selected using the same procedure as for the human cell lines.

**Table 6:** Selected ENCODE tracks for Mouse AID experiment.

| Track | Cell line | ENCODE Accession | Summary |
| --- | --- | --- | --- |
| DNase-seq | ES-e14 | ENCFF672DJH | max |
| H3K36me3 | ES-e14 | ENCFF703RVT | max |
| H3K4me1 | ES-e14 | ENCFF273JBE | max |
| H3K4me3 | ES-Bruce4 | ENCFF611GSQ | max |
| CTCF motif + | - | - | max |
| CTCF motif - | - | - | max |
| painted CTCF UT + | JM8.N4 | GSM5402610 | max ChIP-Seq max motif+ |
| painted CTCF UT - | JM8.N4 | GSM5402610 | max ChIP-Seq max motif- |
| painted CTCF AID + | JM8.N4 | GSM5402616 | max ChIP-Seq max motif+ |
| painted CTCF AID - | JM8.N4 | GSM5402616 | max ChIP-Seq max motif- |

Fast convolution over the selected tracks was formulated as an ordinary least squares (OLS) problem:

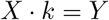

where *X* is a matrix of sliding windows across the genomic track, *k* is the kernel, and *Y* is the target dLEM parameter. The system was regularized using two parameters: *α*, which controls L1 regularization on the kernel, and *λ*, which enforces smoothness of the kernel. To optimize the convolution hyperparameters—including kernel length *k, α, λ*, and a non-negativity constraint on the kernel—we conducted a grid search. The search spanned values of *α* and *λ* from 0 to 1, and tested kernel lengths of 11, 25, 51, 105, and 125.

For each combination of hyperparameters and each track, the optimal kernel was identified using a training set of chromosomes, and performance was assessed on a holdout set (chr19, chr11, chr12) by measuring the correlation with the dLEM parameters in untreated data. The final choice of kernel and hyperparameters was based on the best average correlation to the *L* and *R* parameters, averaged across tracks. This yielded optimal values of *α* = 0, *λ* = 1, kernel length *k* = 51, and a non-negativity constraint set to `True`.

Only tracks from untreated cells were used for linear model fitting. Using a selected set of tracks (DNase, H3K36me3, H3K4me1, H3K4me3, CTCF painted pos/neg, and CTCF motif pos/neg), we trained linear models to predict each of the dLEM parameters for both treated (the ΔCTCF model shown in purple in Fig. 4G, H) and untreated (the UT model, shown in green in Fig. 4G, H) cells. Model performance was evaluated using Pearson correlation between predicted and actual values across all pairwise combinations of two holdout chromosomes. Additionally, we tested the UT models on auxin-treated cell dLEM predictions, and evaluated a modified UT model where all coefficients for CTCF-related tracks were down-weighted by a factor of 0.5. One-sided Wilcoxon test was used to compare the performance of different models.

### Deep dLEM model

Our model consists of two modules: the head and the tail. The head is a U-Net–like convolutional architecture that parameterizes the deep dLEM, while the tail is a flexible module that pools features from sequence data and epigenomic tracks. For the results presented in this paper, we used a simple tail that combines nucleotide-resolution sequence features and DNase-seq signals, applying an attention pooling layer to produce features binned at the resolution of the contact map (10 kb). Instead of providing the model with the full sequence, we supplied only CTCF directional motif matches. When desired, a convolutional tower can also be used as the tail to directly operate on the raw sequence.

The head begins with batch normalization over the combined representation of sequence and epigenomic features. The encoder (downsampling path) is composed of 1D dilated convolutions with kernel size 3 and ReLU activation, followed by depthwise strided convolutions with kernel size 2 and stride 2, also with ReLU activation. Dilated convolutions preserve sequence length while expanding the receptive field, whereas strided convolutions progressively halve the sequence length. Intermediate outputs are stored for skip connections, which later fuse encoder and decoder representations. This design enables the model to capture dependencies across multiple receptive fields and scales of chromatin organization.

The decoder mirrors the encoder structure, progressively upsampling the compressed embedding back to the original sequence length using transposed convolutions. Each upsampling block begins with a strided transposed convolution (kernel size 2, stride 2) followed by ReLU activation, and is paired with a dilated transposed convolution (kernel size 3, dilation 2) also followed by ReLU. At each stage, skip connections inject the corresponding encoder features into the decoder pathway, enriching the reconstruction with multi-scale information. In the implementation used here, both encoder and decoder consist of seven stages, reducing 128 bins at 10 kb resolution down to a single latent unit and then reconstructing them back to 128 bins.

Finally, a 1D convolution with kernel size 1 and two output channels, followed by a sigmoid activation, projects the decoder embedding into the left and right parameter maps required by the deep dLEM. These parameters are then used by the deep dLEM module to generate diagonals of the contact map at any specified index. For training, we employed two supervision strategies: (i) teacher forcing, in which the i-th diagonal is provided to the deep dLEM and the prediction is compared against the (i+1)-th diagonal, and (ii) recursive prediction, in which the i-th diagonal is given and the model is tasked with predicting the diagonal k steps ahead (i+k), with k sampled uniformly between 1 and 15.

The model was trained on 128 *×* 128 contact map patches at 10 kb resolution, derived from all chromosomes except chromosomes 8 and 9, which were reserved for validation and testing, respectively, in the HFF cell line. The H1 cell line was held out entirely for out-of-distribution evaluation. Leveraging the lightweight design of DLEM, which requires only a single diagonal of the contact map to reside in GPU memory at a time, we were able to use a large batch size of 64 on one NVIDIA A4500 GPU. Training was performed on sliding windows of the genome, with consecutive patches overlapping by half the patch size (64 bins), ensuring smoother coverage of genomic loci.

### Comparison with ORCA

We used the 2M ORCA model, the highest-resolution variant, which produces contact maps for 2 Mb genomic regions in a 250 *×* 250 matrix (8 kb resolution). To enable direct comparison with DLEM outputs (128 *×* 128 matrices at 10 kb resolution), we cropped 45 bins from both the start and end of the ORCA predictions and applied the `zoom` function from `scipy.ndimage` to rescale the contact maps to the matching resolution. Because ORCA predictions are not context-aware, we restricted comparisons to chromosome 9, the held-out test split for both models, in the HFF cell line, which is also the training cell line for both methods.

### Comparison with C.Origami

We compared our model to C.origami, trained on IMR90, using predictions on Chromosome 1 of the H1 cell line. As input to C.origami, we used CTCF ChIP-seq and ATAC-seq signals processed with C.origami’s recommended pipeline. C.origami predicts contact maps for 2,097,152 bp windows as 256 *×* 256 matrices, corresponding to an 8192 bp bin size. To align with DLEM, we matched patch starts and extracted the first 157 bins of each C.origami patch, spanning 1,286,144 bp, comparable to the 1,280,000 bp covered by DLEM. Finally, we applied `zoom` to rescale the 8192 bp resolution of C.origami to the 8000 bp resolution of DLEM.

### Code availability

Source code and analysis scripts for directly fitting dLEM to contact map data are available at https://github.com/chikinalab/dLEM. Source code for deep dLEM is available at https://github.com/chikinalab/dLEMpytorch. dLEM is distributed as a Python package for easy installation and use.

### Derivation of dLEM

The differentiable loop extrusion model (dLEM) provides an interpretable mechanistic description of how cohesin-mediated loop extrusion shapes chromatin contact patterns observed in chromosome conformation capture experiments. By formulating loop extrusion as a differentiable process (one in which the relationship between model parameters and output can be expressed through smooth continuous functions that allow gradient-based optimization), dLEM bridges the gap between physical mechanisms and data-driven inference, enabling both mechanistic understanding and efficient parameter estimation. Our derivation begins from the first principles of the loop extrusion process and makes several key simplifying assumptions that enable tractable, genome-wide inference while retaining the essential physics.

In the loop extrusion process, cohesin complexes bind to chromatin and translocate bidirectionally along the fiber. Each cohesin can be characterized by the genomic positions of its two contact points: position *i* for the left anchor and position *j* for the right anchor. We model the dynamics of cohesin translocation as a continuous-time Markov process on the two-dimensional lattice of possible contact positions (*i, j*). A key modeling assumption underlying dLEM is that the population-averaged contact map can be approximated as the steady-state distribution of these cohesin positions. This represents an idealization, as chromsome conformation capture experiments measure contacts between DNA fragments rather than direct cohesin positions, other factors contribute to contact formation, and the population-averaged conformation is likely not physically realizable in individual cells. Nevertheless, this approximation captures the dominant contribution of loop extrusion to chromatin architecture and provides a tractable framework for mechanistic inference. With this assumption in mind, consider *C*^*i,j*^ as the expected number of cohesin complexes bridging positions *i* and *j*. By neglecting potential traffic jams or cooperative effects between multiple cohesins, the temporal evolution of this quantity is governed by the following master equation:

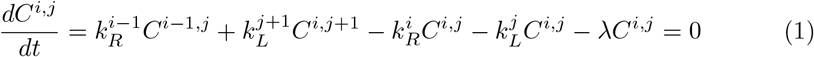

where:

- 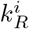 represents the transition rate for rightward movement from position *i* to *i* + 1
- 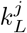 represents the transition rate for leftward movement from position *j* to *j −* 1
- *λ* is the constant detachment rate at which cohesin dissociates from chromatin (while cohesin release is likely locally modulated, a constant *λ* provides an approximation that can be robustly estimated from distance-decay curves).

The first two terms represent influx: cohesin flowing into state (*i, j*) from neighboring states through rightward movement from (*i−* 1, *j*) and leftward movement from (*i, j* + 1). The next two terms represent outflux through continued translocation, and the final term represents loss through detachment. At steady state, these fluxes balance, yielding:

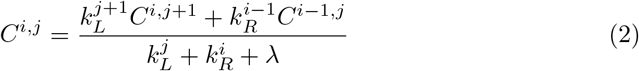

The transition rates *k*_*R*_ and *k*_*L*_ must be derived from the underlying physical process of cohesin translocation. Here, we introduce another key assumption: the left and right legs of cohesin move independently, with position-specific velocities that can be modulated by chromatin-bound factors. This independence assumption, while a simplification of the coupled mechanical system, captures the primary effect that impediments (such as CTCF) affect each leg separately depending on their orientation. Under this independence assumption, we can model each leg’s dynamics separately. For a single leg at position *x* with velocity *v*_leg_(*x*), the concentration *C*_leg_(*x, t*) evolves according to the continuity equation:

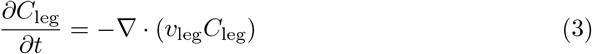

This continuous description must be discretized to match the resolution of chromosome conformation capture experiments, where the genome is divided into bins of size *h*. Loop extrusion is fundamentally an advective (directed transport) process: cohesin complexes move with a definite direction and velocity along the chromatin fiber. To discretize the equation, we can use the upwind finite difference scheme:

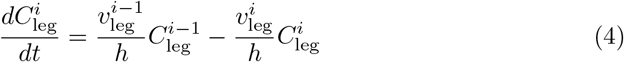

This discretization has a clear physical interpretation: the factor *v/h* represents the rate at which cohesin exits a bin of size *h* when moving with velocity *v*.

Finally, to create a scale-free parameterization suitable for optimization, we factor out the maximum translocation velocity *v*_max_ and express velocities as normalized profiles:

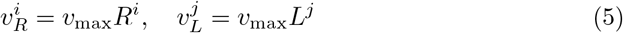

where *R*^*i*^, *L*^*j*^ ∈ [0, 1] represent the fraction of maximum velocity at each position. This yields transition rates:

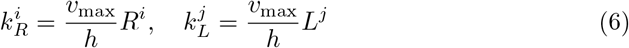

Incorporating the scale-free velocity parameterization, the detachment rate becomes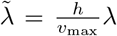 (notice that the same fraction occurs in the distance-decay model as an exponential decay parameter), yielding the final expression for the steady state number of cohesins bound to loci *i* and *j*:

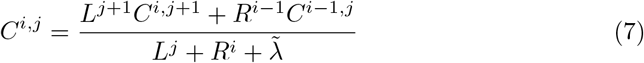

This equation describes how the contact frequency at each pixel emerges from the balance between cohesin influx from neighboring contacts and outflux through translocation and detachment.

The dLEM formulation makes several simplifying assumptions: independent leg movement, steady-state approximation, constant detachment rate, and neglect of cohesin-cohesin interactions. Additionally, dLEM focuses exclusively on loop extrusion dynamics omitting other mechanisms of genome organization (e.g. compartmentalization and interaction with the nuclear lamina). This choice is motivated by the observation that at the length scale of several hundred kb, loop extrusion creates the characteristic differential patterns visible in contact maps (TADs, stripes, loops), while polymer crumpling contribution can be approximated with and the overall distance-decay background that can be approximated as an independent component and added back separately (see main text, Fig. 1E). These simplifications reflect a deliberate modeling choice prioritizing interpretability and computational tractability, and dLEM successfully captures the major local features of contact maps (Fig. 2) while providing an interpretable, one-dimensional description of loop extrusion dynamics that can be directly compared with other genomic features.

### Distance-decay model for constraining detachment rate

While dLEM successfully captures local contact patterns through position-specific velocity profiles, the global detachment rate *λ* remains poorly constrained when fitting to contact maps alone, creating an identifiability problem during optimization. To address this, we developed a complementary distance-decay analysis that describes average genome-wide contact frequency patterns using a minimal sufficient model for contact decay:

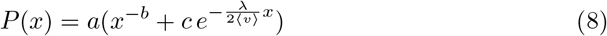

We model the average contact frequency *P* (*x*) as a function of genomic distance *x* as arising from two independent contributions: passive polymer folding, which produces a power-law decay *x*^*−b*^, and active loop extrusion, which generates an exponential decay component *e*^*−λ/*(2⟨*v*⟩)*x*^ where ⟨*v*⟩ represents the average cohesin velocity.

Rather than imposing the theoretical *−*3*/*2 scaling characteristic of a Gaussian chain, we treat the power-law exponent *b* as a free parameter to account for effects such as nuclear packing and loop extrusion itself [25]. The exponential decay component arises from our assumption that cohesin complexes have a constant probability of detachment per unit time, yielding exponential decay kinetics when averaged over all genomic fragments of length *x*. While individual cohesin complexes experience heterogeneous detachment rates due to factors like WAPL co-localization with CTCF, this population-level approximation captures the overall statistical behavior. The length scale emerges from the time scale through the velocity parameter ⟨*v*⟩, which similarly represents a population average despite local variations in cohesin translocation rates.

While polymer folding and loop extrusion likely exhibit some interdependence, we treat them as additive contributions in this minimal model. We assume loop extrusion dynamics remain largely independent of polymer folding, and even if the underlying parameters are correlated, the overall contact frequency patterns should still exhibit separable power-law and exponential components. The model produces quantitatively accurate fits to experimental distance-decay data across all human chromosomes (Fig. S1), supporting the sufficiency of this model for a first-order approximation. The parameter *c* indicates the relative contribution of loop extrusion to overall contact frequency, while *a* provides data scaling. By fitting the composite model to genome-wide distance-decay curves, we reliably estimate *λ* independently of local velocity profiles, providing the constraint necessary for robust dLEM parameter estimation.

## Supplementary Figures

**Fig. S1:**
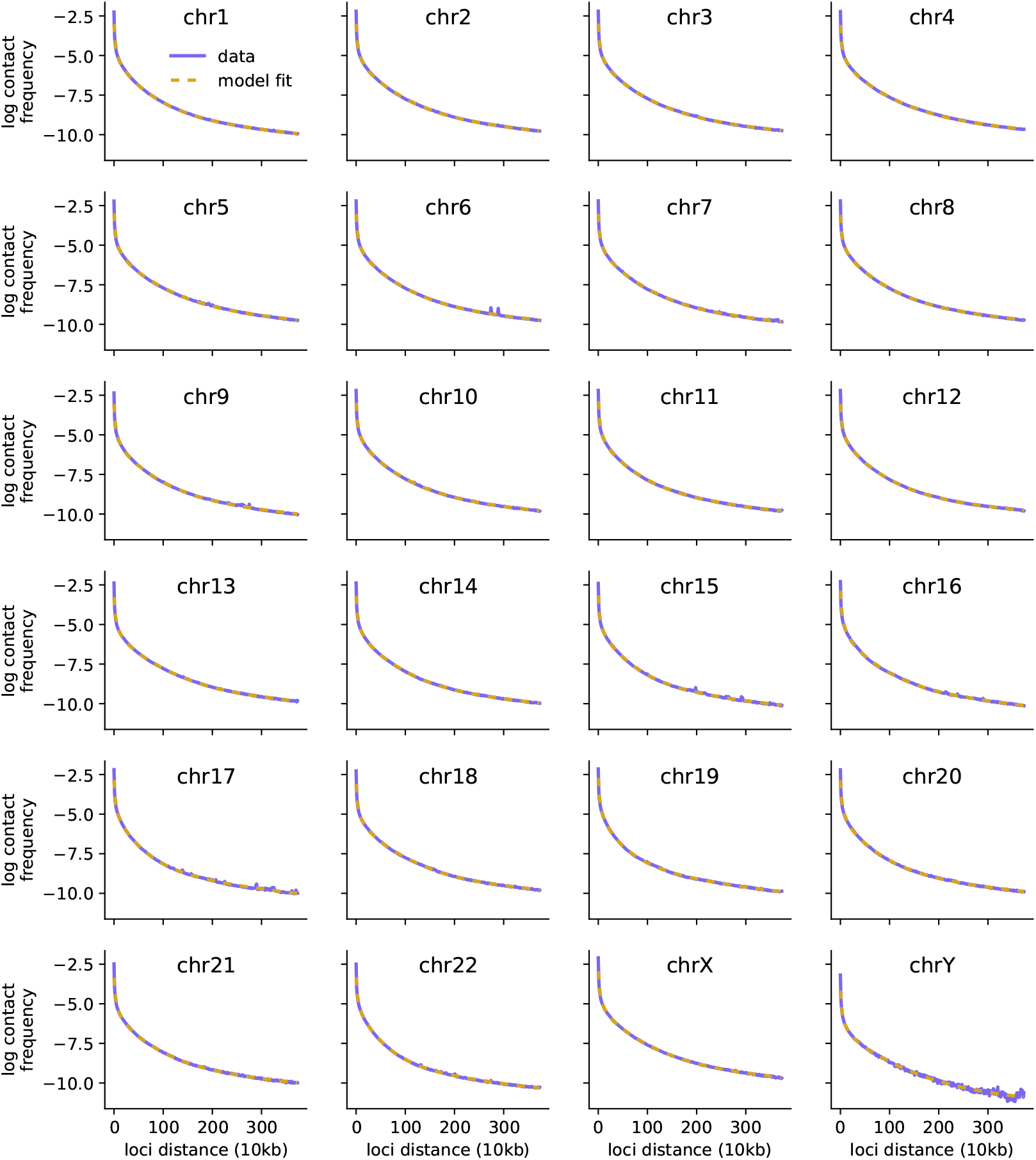
Distance-decay analysis constrains cohesin detachment rate across all human chromosomes. Contact frequency decay as a function of genomic distance for all 24 human chromosomes in H1hESC cells. Solid violet lines show experimental average contact frequencies (log scale) for loci separated by 10kb to 4Mb, binned at 10kb resolution. Golden dashed lines show fits of the diffusion-extrusion model, which decomposes distance-dependent contact decay into passive polymer diffusion (power-law component) and active loop extrusion (exponential component). Consistent fits across all chromosomes validate the model’s ability to constrain the global detachment rate parameter *λ* used in dLEM fitting.

**Fig. S2:**
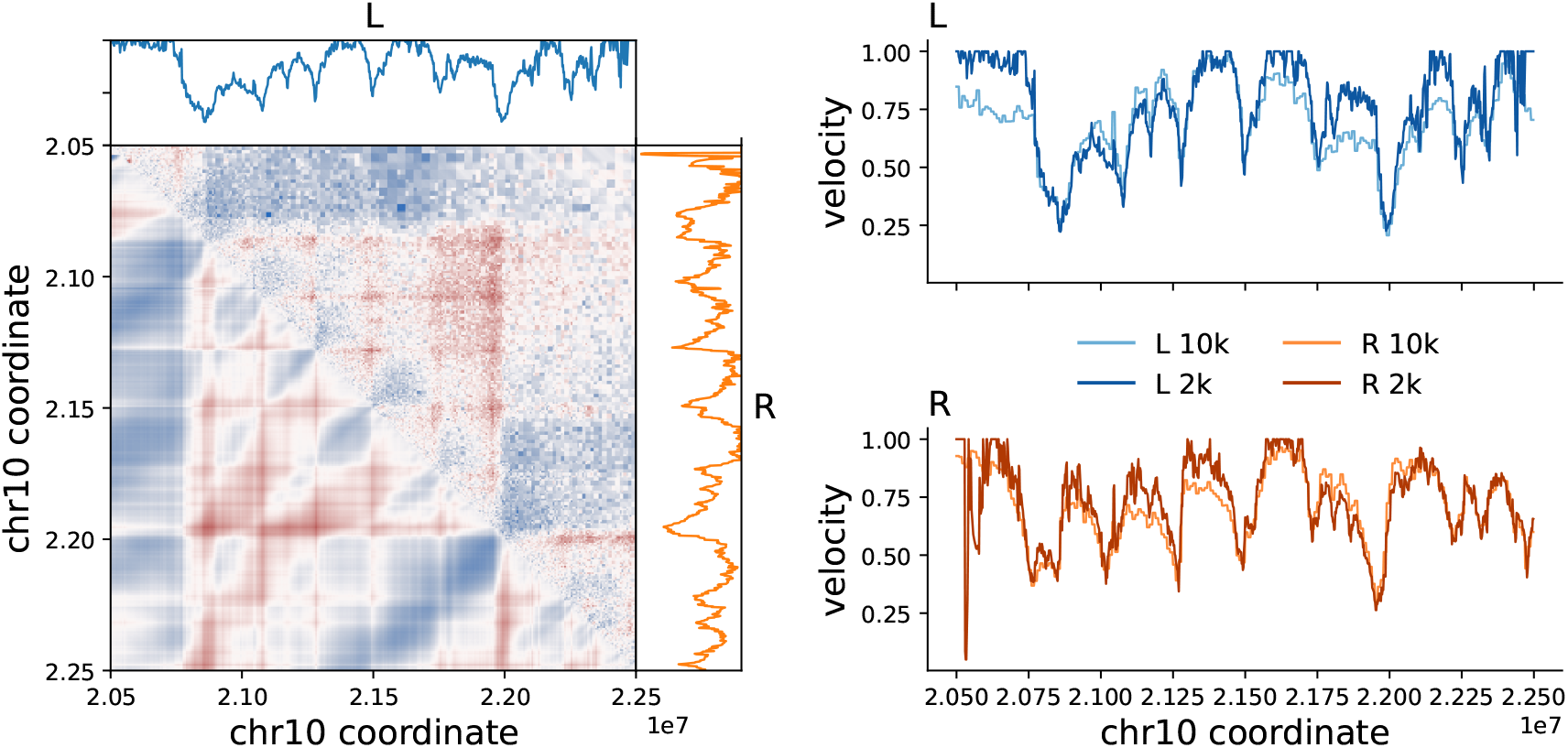
dLEM consistency across resolutions. Same genomic region as Fig. 2 A-B (chr10:20.5-22.5Mb), showing data and model fit at 2kb resolution. Velocity parameter comparison (right panels) demonstrates high correspondence between 10kb (light colors) and 2kb (dark colors) fitted parameters for both L (blue) and R (orange), indicating robust parameter estimation across different resolutions.

**Fig. S3:**
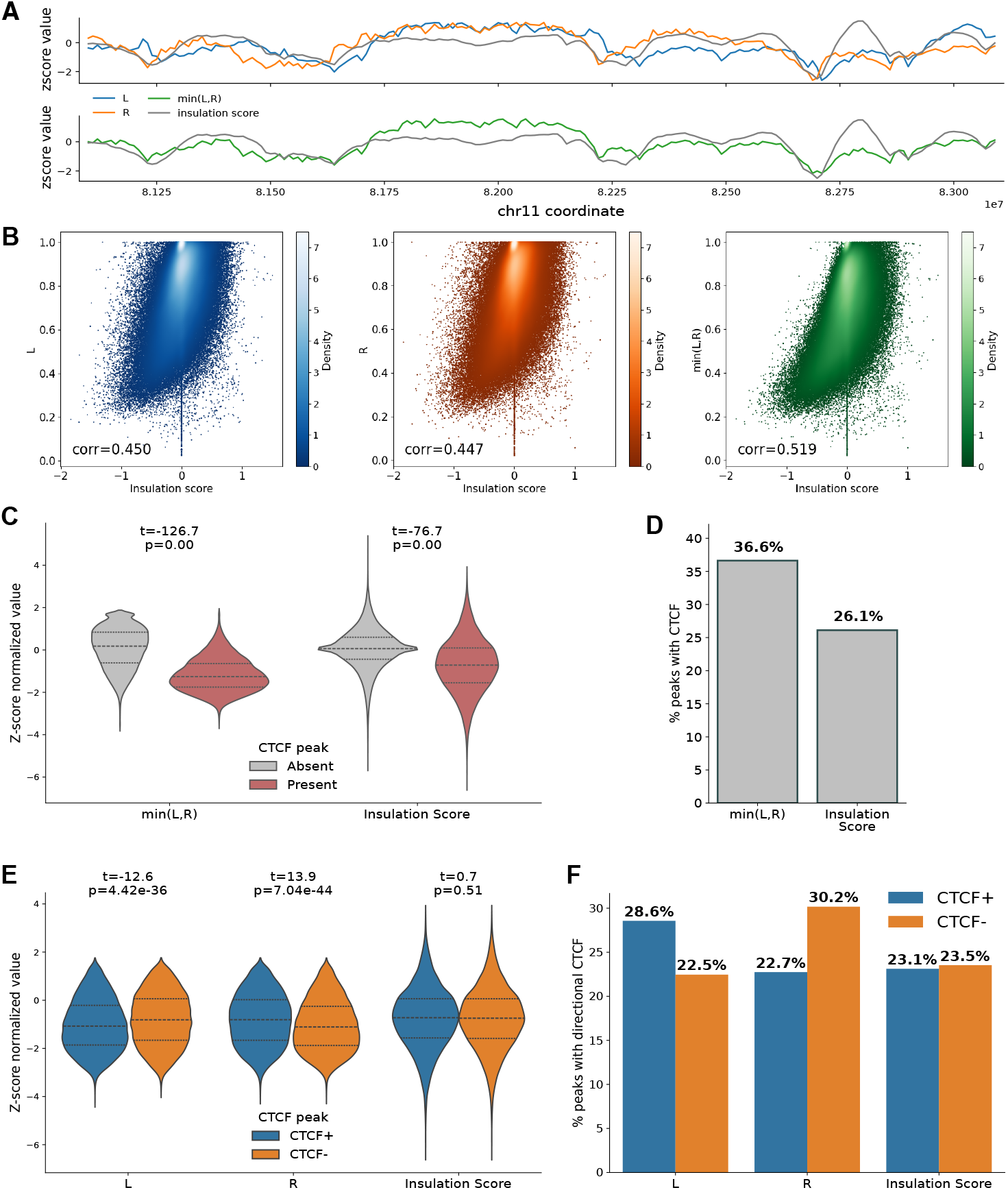
**A**: *dLEM parameters correspond with insulation scores*. Example genomic region (chr11:81100000-83100000) showing *L, R*, min(*L, R*), and diamond insulation scores (z-score normalized values). **B**: *dLEM parameters correlate with insulation scores*. Scatter plots show correlation between insulation scores and *L* (corr=0.450), *R* (corr=0.447), and min(*L, R*) (corr=0.519), with min(*L, R*) achieving the highest correlation. **C**: min(*L, R*) *separates CTCF peak distributions more strongly than insulation scores*. Violin plots comparing z-score distributions at regions with CTCF peaks present versus absent show that min(*L, R*) achieves stronger separation (t=-126.7, p=0.00) than insulation scores (t=-76.7, p=0.00). **D**: min(*L, R*) *minima are enriched for CTCF peaks*. 36.6% of min(*L, R*) minima (*z* ≤ *−*2) contain CTCF peaks compared to 26.1% of insulation score minima. **E**: *L and R separate CTCF orientation*. Violin plots show *L* is significantly lower at CTCF+ peaks (t=-12.6, p=4.42e-36) while *R* is significantly lower at CTCF-peaks (t=13.9, p=7.04e-44). Insulation scores show no directional preference (t=0.7, p=0.51). **F**: *L and R minima are enriched for convergent CTCF. L* minima contain 28.6% CTCF+ versus 22.5% CTCF-peaks, and *R* minima contain 30.2% CTCF-versus 23.5% CTCF+ peaks. Insulation score minima show similar proportions (22.7% CTCF+ versus 23.1% CTCF-). All analyses: 10kb resolution, chromosomes 3-8 and 10-13, H1-hESC.

**Fig. S4:**
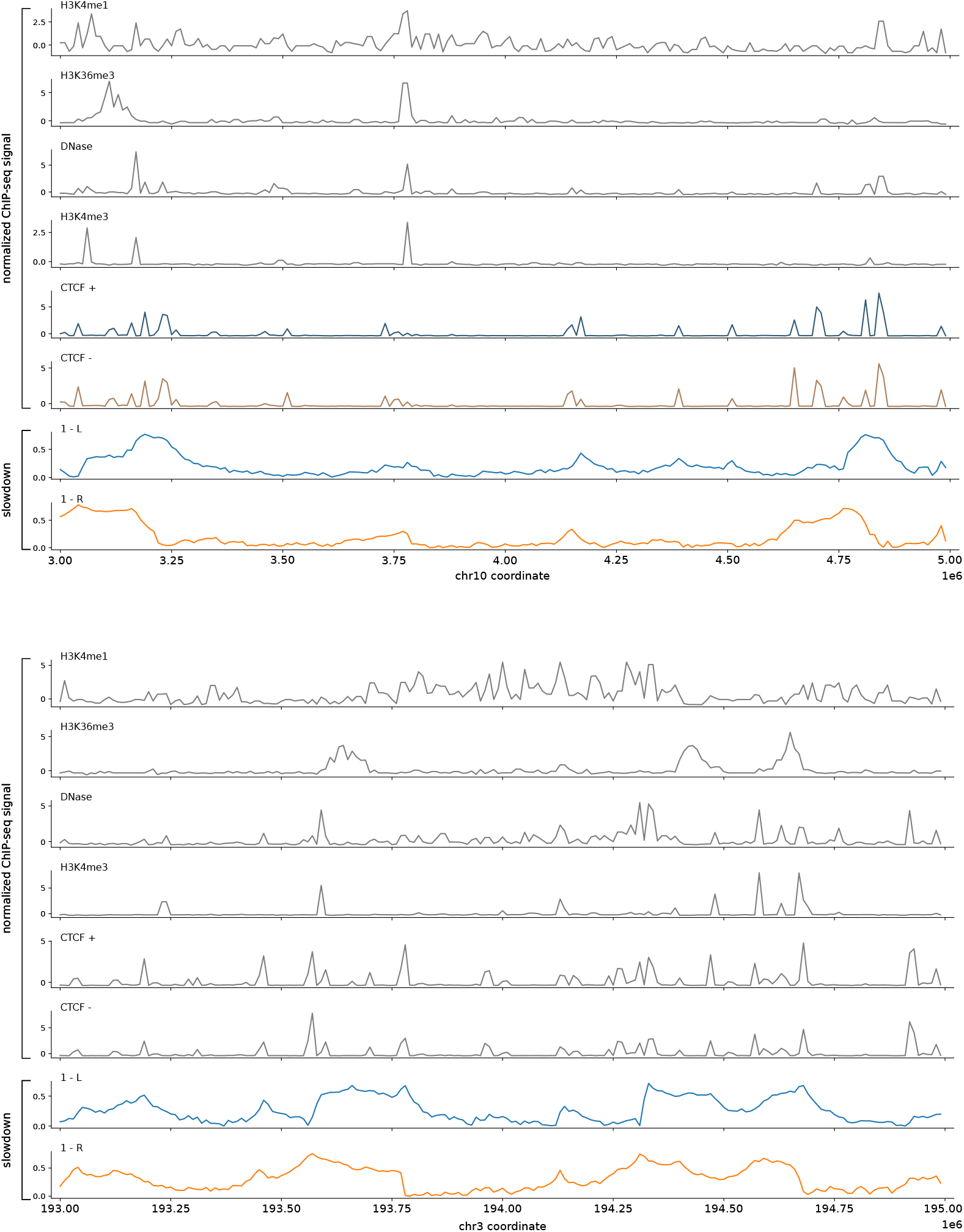
Comparison of genomic tracks and slowdown profiles (1 − *L* and 1 − *R*) for two genomic regions (chr10:3000000-5000000 and chr3:193000000-195000000) in H1hESC in 10kb resolution.

**Fig. S5:**
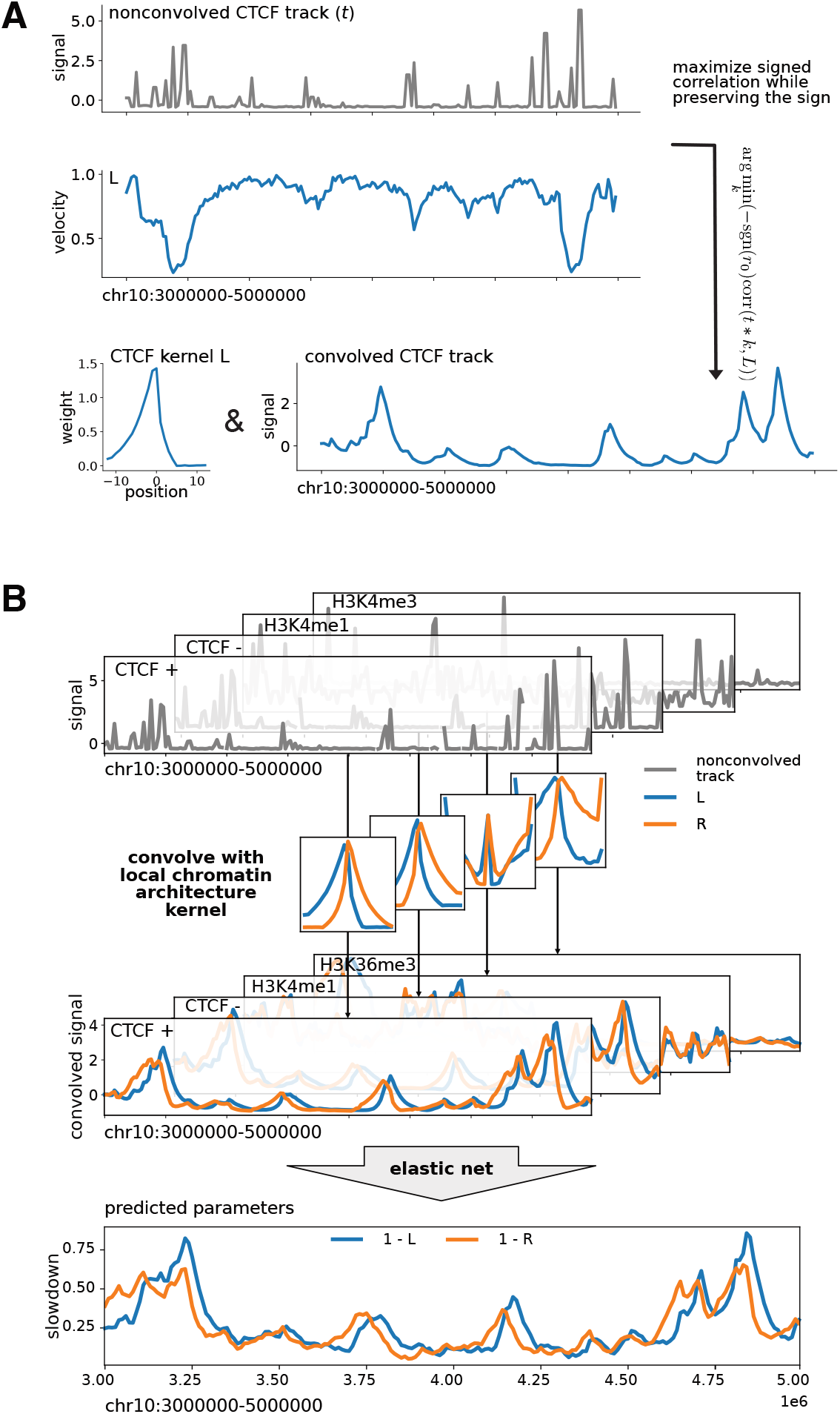
**A**: *Pipeline for learning local chromatin architecture kernels* Kernels are optimized to maximize signed correlations between individual genomic tracks and dLEM parameters while preserving the original sign of the correlations. **B**: *Linear model pipeline for predicting dLEM parameters from chromatin features*. ENCODE ChIP-seq tracks (top) are binned to contact map resolution (10kb) and convolved with parameter-specific optimized kernels to capture local chromatin architecture effects, then used in elastic net regression to predict L and R velocity parameters. CTCF features are directionally signed using motif orientation to distinguish forward and reverse binding effects.

**Fig. S6:**
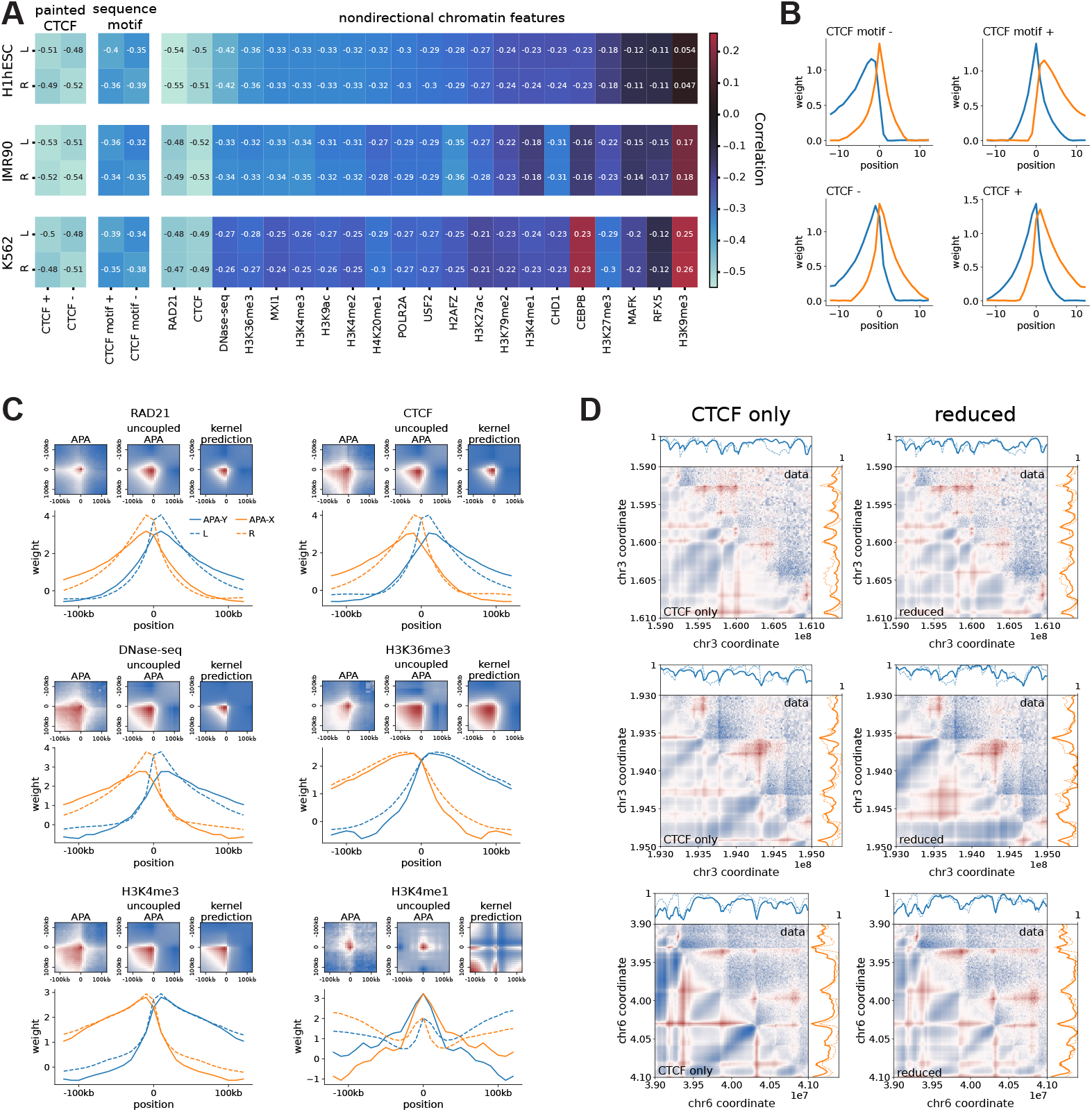
**A**: *Convolution with local chromatin architecture kernels enhances feature-parameter correlations* Comparison of correlations between dLEM parameters and genomic features for H1hESC, IMR90, and K562 cell lines (chr3-8 and chr10-13) after applying convolution kernels. Convolution consistently improves correlations while preserving directional asymmetries in CTCF. **B**: *Learned kernels of directional CTCF features retain orientation-specific asymmetry* Optimized convolution kernels for CTCF motif orientation and directionally-signed ChIP-seq tracks show distinct asymmetric profiles, consistent with the known directional blocking effects of CTCF on cohesin translocation. **C**: *Learned kernels recapitulate aggregate peak analysis (APA) patterns* Comparison of learned convolution kernels with APA plot marginal distributions demonstrates strong correspondence, indicating that the kernels capture biologically meaningful local chromatin architecture effects. The outer products of *L* and *R* kernels (kernel prediction) resemble both experimental APA plots and their uncoupled approximations (outer product of the APA marginals), validating the mechanistic basis of the convolution approach. **D**: *dLEM predictions from chromatin features*. Three genomic regions showing experimental data (upper triangles) and contact maps generated using *L* and *R* parameters predicted from the either the CTCF-only or reduced-feature ElasticNet model (lower triangles), showing that addition of transcription-related tracks improves the prediction.

**Fig. S7:**
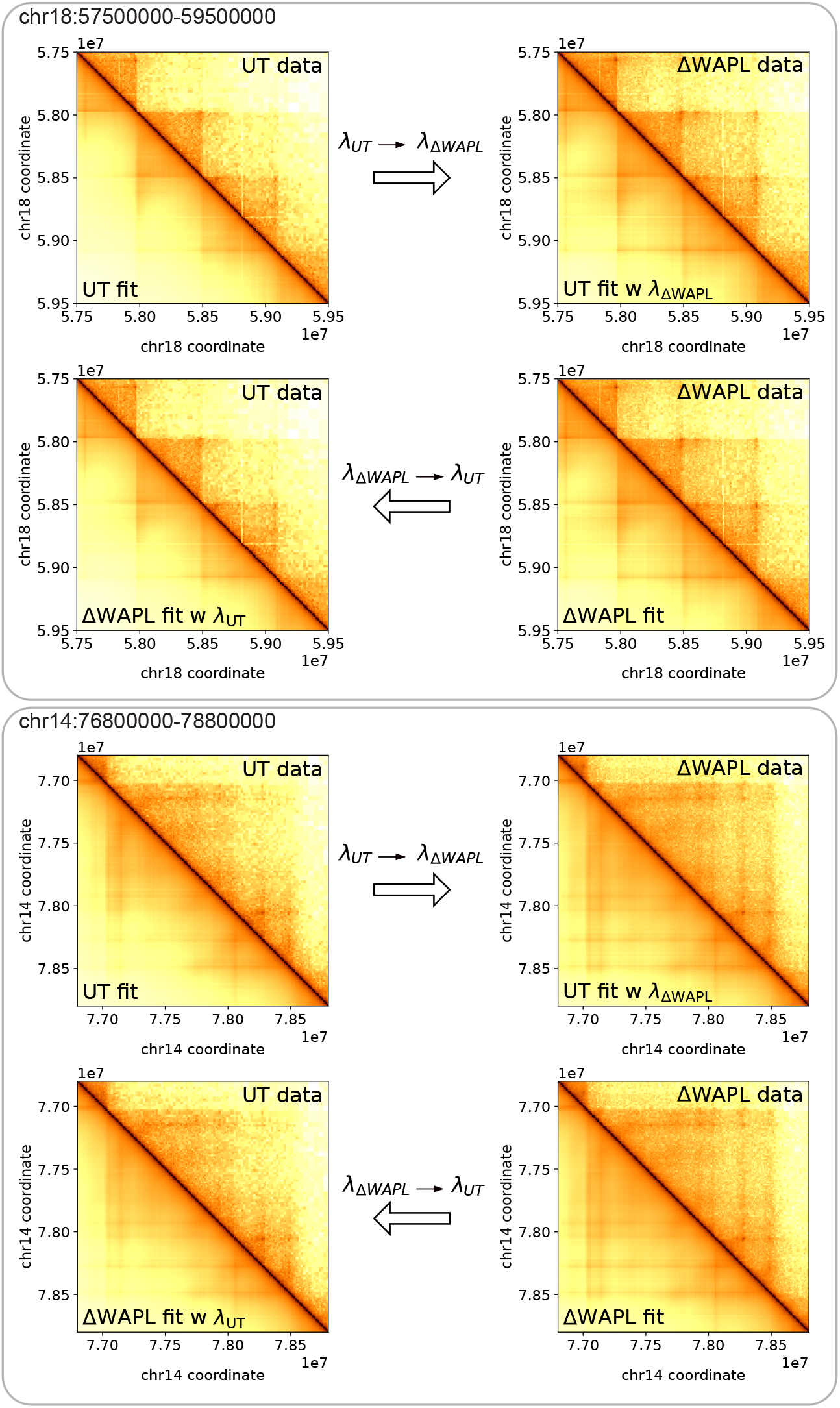
Additional patches showing *λ* parameter successfully captures the effects of ΔWAPL perturbation.

**Fig. S8:**
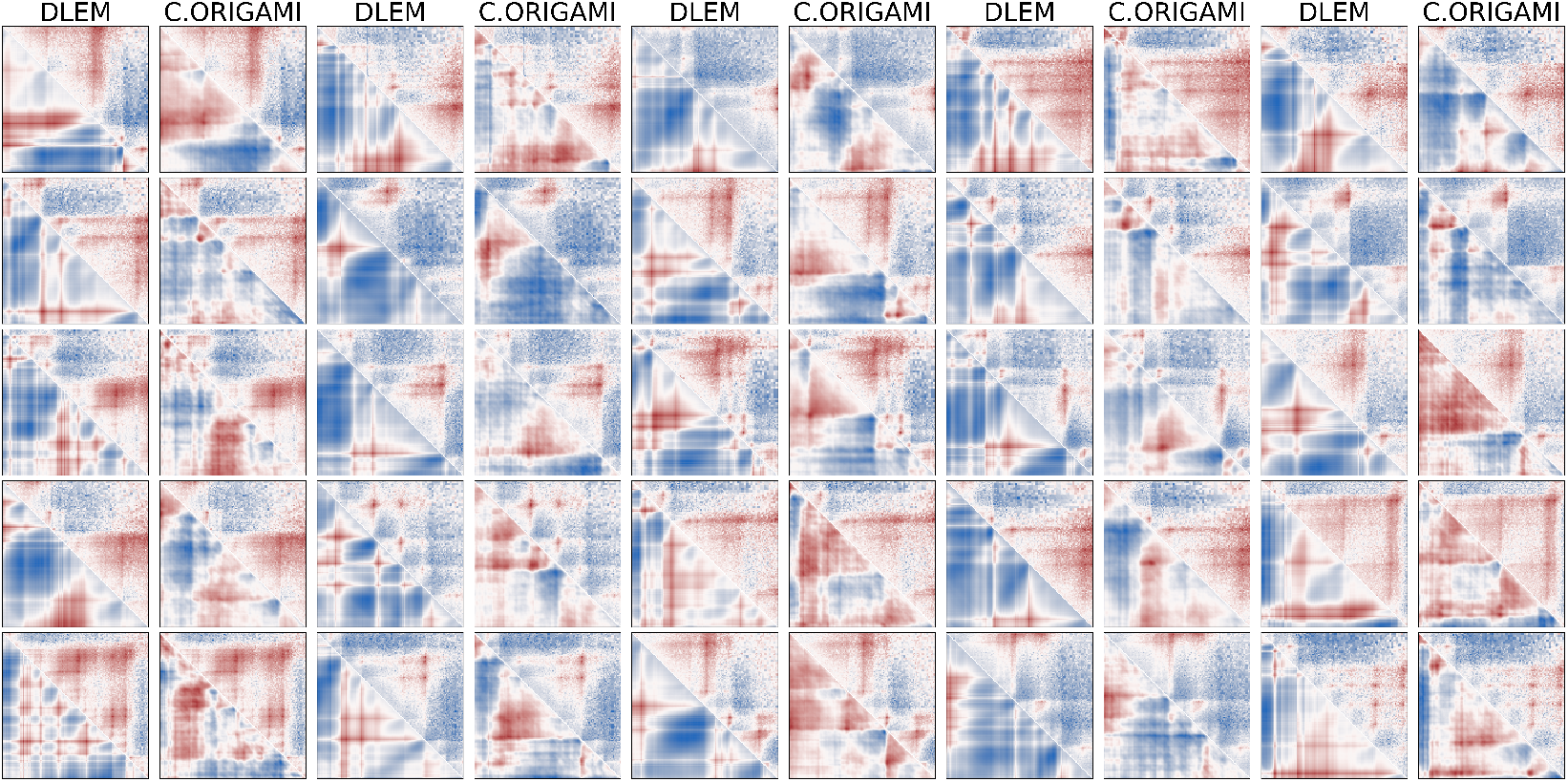
Additional patches comparing C. Origami and deep dLEM.

**Fig. S9:**
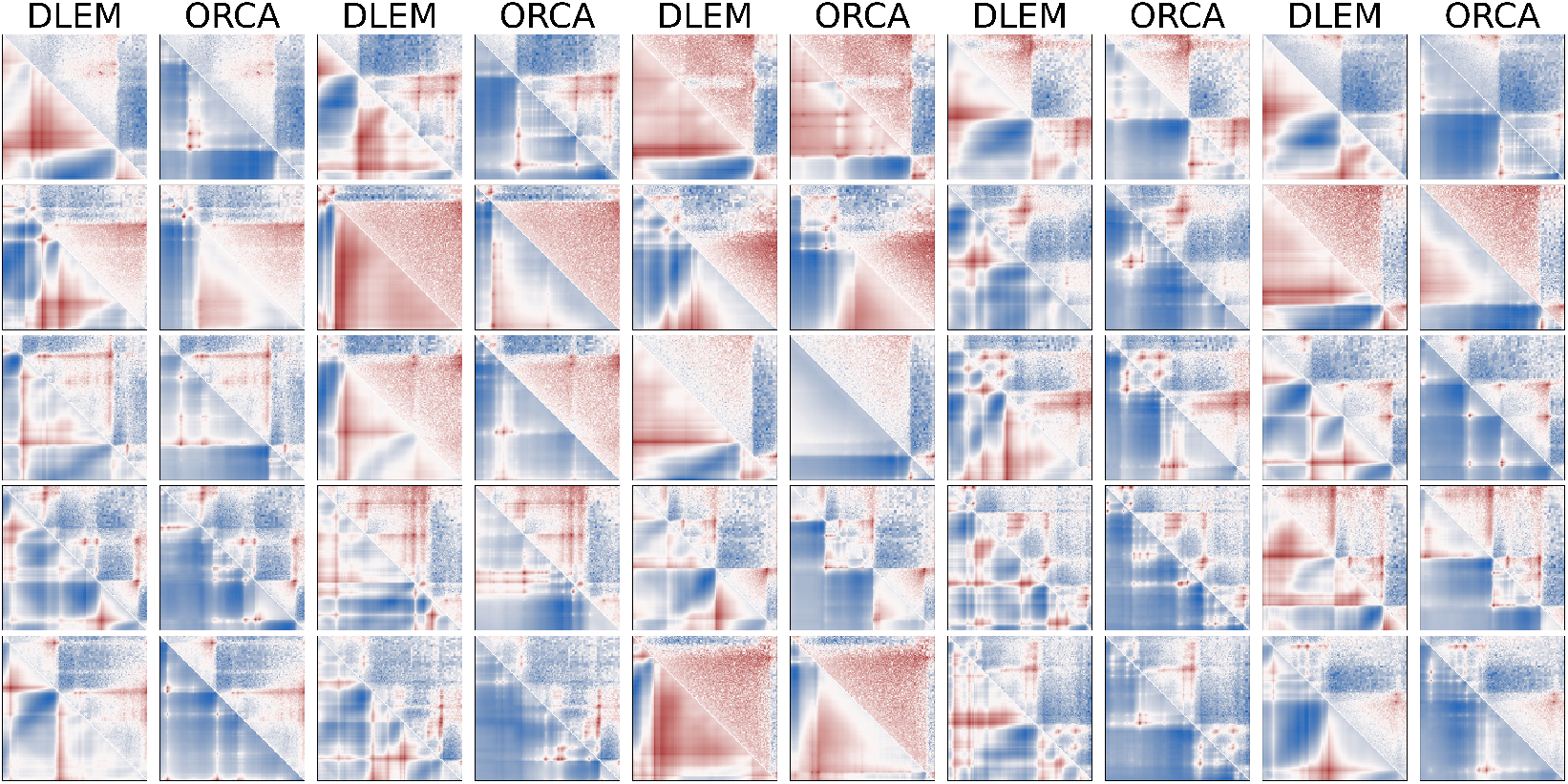
Additional patches comparing Orca and deep dLEM.

## References

[1] Chakraborty, A., Ay, F.: The role of 3d genome organization in disease: From compartments to single nucleotides. Seminars in Cell Developmental Biology 90, 104–113 (2019) 10.1016/j.semcdb.2018.07.005. 3D Genome and Diseases

[2] McCord, R.P., Kaplan, N., Giorgetti, L.: Chromosome conformation capture and beyond: Toward an integrative view of chromosome structure and function. Molecular Cell 77(4), 688–708 (2020) 10.1016/j.molcel.2019.12.021

[3] Rao, S.S.P., Huntley, M.H., Durand, N.C., Stamenova, E.K., Bochkov, I.D., Robinson, J.T., Sanborn, A.L., Machol, I., Omer, A.D., Lander, E.S., Aiden, E.L.: A 3d map of the human genome at kilobase resolution reveals principles of chromatin looping. Cell 159(7), 1665–1680 (2014) 10.1016/j.cell.2014.11.021

[4] Hsieh, T.-H.S., Weiner, A., Lajoie, B., Dekker, J., Friedman, N., Rando, O.J.: Micro-c xl: assaying chromosome conformation from the nucleosome to the entire genome. Nature Methods 17(8), 782–788 (2020)

[5] Dixon, J.R., Selvaraj, S., Yue, F., Kim, A., Li, Y., Shen, Y., Hu, M., Liu, J.S., Ren, B.: Topological domains in mammalian genomes identified by analysis of chromatin interactions. Nature 485(7398), 376–380 (2012) 10.1038/nature11082

[6] Lieberman-Aiden, E., Berkum, N.L., Williams, L., Imakaev, M., Ragoczy, T., Telling, A., Amit, I., Lajoie, B.R., Sabo, P.J., Dorschner, M.O., Sandstrom, R., Bernstein, B., Bender, M.A., Groudine, M., Gnirke, A., Stamatoyannopoulos, J., Mirny, L.A., Lander, E.S., Dekker, J.: Comprehensive mapping of long-range interactions reveals folding principles of the human genome. Science 326(5950), 289–293 (2009) 10.1126/science.1181369

[7] Dekker, J., Mirny, L.A.: The chromosome folding problem and how cells solve it. Cell 187(23), 6424–6450 (2024) 10.1016/j.cell.2024.10.026

[8] Fudenberg, G., Imakaev, M., Lu, C., Goloborodko, A., Abdennur, N., Mirny, L.A.: Formation of chromosomal domains by loop extrusion. Cell Reports 15(9), 2038–2049 (2016) 10.1016/j.celrep.2016.04.085

[9] Nuebler, J., Fudenberg, G., Imakaev, M., Abdennur, N., Mirny, L.A.: Chromatin organization by an interplay of loop extrusion and compartmental segregation. Proceedings of the National Academy of Sciences 115(29), 6697–6706 (2018) 10.1073/pnas.1717730115

[10] Stigler, J., Çamdere, G., Koshland, D.E., Greene, E.C.: Single-molecule imaging reveals a collapsed conformational state for dna-bound cohesin. Cell Reports 15(5), 988–998 (2016) 10.1016/j.celrep.2016.04.003

[11] Kim, Y., Shi, Z., Zhang, H., Finkelstein, I.J., Yu, H.: Human cohesin compacts dna by loop extrusion. Science 366(6471), 1345–1349 (2019) 10.1126/science.aaz4475

[12] Heinz, S., Texari, L., Hayes, M.G.B., Urbanowski, M., Chang, M.W., Givarkes, N., Rialdi, A., White, K.M., Albrecht, R.A., Pache, L., Marazzi, I., García-Sastre, A., Shaw, M.L., Benner, C.: Transcription elongation can affect genome 3d structure. Cell 174(6), 1522–153622 (2018) 10.1016/j.cell.2018.07.047

[13] Valton, A.-L., Venev, S.V., Mair, B., Khokhar, E.S., Tong, A.H.Y., Usaj, M., Chan, K., Pai, A.A., Moffat, J., Dekker, J.: A cohesin traffic pattern genetically linked to gene regulation. Nature Structural & Molecular Biology 29(12), 1239–1251 (2022)

[14] Banigan, E.J., Tang, W., Berg, A.A., Stocsits, R.R., Wutz, G., Brandão, H.B., Busslinger, G.A., Peters, J.-M., Mirny, L.A.: Transcription shapes 3d chromatin organization by interacting with loop extrusion. Proceedings of the National Academy of Sciences 120(11), 2210480120 (2023) 10.1073/pnas.2210480120

[15] Rao, S.S.P., Huang, S.-C., St Hilaire, B.G., Engreitz, J.M., Perez, E.M., Kieffer-Kwon, K.-R., Sanborn, A.L., Johnstone, S.E., Bascom, G.D., Bochkov, I.D., Huang, X., Shamim, M.S., Shin, J., Turner, D., Ye, Z., Omer, A.D., Robinson, J.T., Schlick, T., Bernstein, B.E., Casellas, R., Lander, E.S., Aiden, E.L.: Cohesin loss eliminates all loop domains. Cell 171(2), 305–32024 (2017) 10.1016/j.cell.2017.09.026

[16] Zuin, J., Roth, G., Zhan, Y., Cramard, J., Redolfi, J., Piskadlo, E., Mach, P., Kryzhanovska, M., Tihanyi, G., Kohler, H., Eder, M., Leemans, C., Steensel, B., Meister, P., Smallwood, S., Giorgetti, L.: Nonlinear control of transcription through enhancer–promoter interactions. Nature 604(7906), 571–577 (2022) 10.1038/s41586-022-04570-y

[17] Crane, E., Bian, Q., McCord, R.P., Lajoie, B.R., Wheeler, B.S., Ralston, E.J., Uzawa, S., Dekker, J., Meyer, B.J.: Condensin-driven remodelling of x chromosome topology during dosage compensation. Nature 523(7559), 240–244 (2015) 10.1038/nature14450

[18] Gjoni, K., Gunsalus, L.M., Kuang, S., McArthur, E., Pittman, M., Capra, J.A., Pollard, K.S.: Comparing chromatin contact maps at scale: methods and insights. Nature Methods 22(4), 824–833 (2025) 10.1038/s41592-025-02630-5

[19] Fu, Y., Zhao, T., Clark, F., Nomikou, S., Tsirigos, A., Lionnet, T.: Connecting chromatin structures to gene regulation using dynamic polymer simulations. eLife (2024) 10.7554/elife.94738.2

[20] Schwessinger, R., Gosden, M., Downes, D., Brown, R.C., Oudelaar, A.M., Telenius, J., Teh, Y.W., Lunter, G., Hughes, J.R.: Deepc: predicting 3d genome folding using megabase-scale transfer learning. Nature Methods 17(11), 1118–1124 (2020) 10.1038/s41592-020-0960-3

[21] Fudenberg, G., Kelley, D.R., Pollard, K.S.: Predicting 3d genome folding from dna sequence with akita. Nature Methods 17(11), 1111–1117 (2020) 10.1038/s41592-020-0958-x

[22] Zhou, J.: Sequence-based modeling of three-dimensional genome architecture from kilobase to chromosome scale. Nature Genetics 54(5), 725–734 (2022) 10.1038/s41588-022-01065-4

[23] Tan, J., Shenker-Tauris, N., Rodriguez-Hernaez, J., Wang, E., Sakellaropoulos, T., Boccalatte, F., Thandapani, P., Skok, J., Aifantis, I., Fenyö, D., Xia, B., Tsirigos, A.: Cell-type-specific prediction of 3d chromatin organization enables high-throughput in silico genetic screening. Nature Biotechnology 41(8), 1140–1150 (2023) 10.1038/s41587-022-01612-8

[24] Smaruj, P.N., Xiao, Y., Fudenberg, G.: Recipes and ingredients for deep learning models of 3d genome folding. Current Opinion in Genetics Development 91, 102308 (2025) 10.1016/j.gde.2024.102308

[25] Polovnikov, K.E., Brandão, H.B., Belan, S., Slavov, B., Imakaev, M., Mirny, L.A.: Crumpled polymer with loops recapitulates key features of chromosome organization. Phys. Rev. X 13, 041029 (2023) 10.1103/PhysRevX.13.041029

[26] Hsieh, T.-H.S., Cattoglio, C., Slobodyanyuk, E., Hansen, A.S., Darzacq, X., Tjian, R.: Enhancer–promoter interactions and transcription are largely maintained upon acute loss of ctcf, cohesin, wapl or yy1. Nature Genetics 54(12), 1919–1932 (2022) 10.1038/s41588-022-01223-8

[27] Haarhuis, J.H.I., van der Weide, R.H., Blomen, V.A., Yáñez-Cuna, J.O., Amendola, M., van Ruiten, M.S., Krijger, P.H.L., Teunissen, H., Medema, R.H., van Steensel, B., Brummelkamp, T.R., de Wit, E., Rowland, B.D.: The cohesin release factor wapl restricts chromatin loop extension. Cell 169(4), 693–70714 (2017) 10.1016/j.cell.2017.04.013

[28] Karczewski, K.J., Francioli, L.C., Tiao, G., Cummings, B.B., Alföldi, J., Wang, Q., Collins, R.L., Laricchia, K.M., Ganna, A., Birnbaum, D.P., et al.: The mutational constraint spectrum quantified from variation in 141,456 humans. Nature 581(7809), 434–443 (2020) 10.1038/s41586-020-2308-7

[29] Open2C, Abdennur, N., Abraham, S., Fudenberg, G., Flyamer, I.M., Galitsyna, A.A., Goloborodko, A., Imakaev, M., Oksuz, B.A., Venev, S.V., Xiao, Y.: Cooltools: Enabling high-resolution hi-c analysis in python. PLOS Computational Biology 20(5), 1–16 (2024) 10.1371/journal.pcbi.1012067

[30] Paszke, A., Gross, S., Massa, F., Lerer, A., Bradbury, J., Chanan, G., Killeen, T., Lin, Z., Gimelshein, N., Antiga, L., Desmaison, A., Köpf, A., Yang, E.Z., DeVito, Z., Raison, M., Tejani, A., Chilamkurthy, S., Steiner, B., Fang, L., Bai, J., Chintala, S.: Pytorch: An imperative style, high-performance deep learning library. In: Wallach, H.M., Larochelle, H., Beygelzimer, A., Buc, F., Fox, E.B., Garnett, R. (eds.) NeurIPS, pp. 8024–8035 (2019)

